# Structure-guided design of native-like HIV Env Single Chain Trimers for enhanced stability, immunogenicity, and versatile vaccine delivery

**DOI:** 10.1101/2025.10.25.684215

**Authors:** Xuduo Li, Ipsita Krishnamurthy, Nitesh Mishra, Nicole E. James, Rohan Roy Chowdhury, Sean Callaghan, Anjali Somanathan, Gabriel Avillion, Khaled Amereh, Tharunika Sekar, Prabhgun Oberoi, Yuxin Zhu, Jonathan L. Torres, Wen-Hsin Lee, Joel D. Allen, Bo Liang, Gabriella Giese, Anh L. Vo, Sharaf Andrabi, Ishita Jain, Asanti Mergiya, Tazio Capozzola, Wan-ting He, Fabio Anzanello, Rumi Habib, Frederic Bibollet-Ruche, Rebecca Nedellec, Sara Dutton Sackett, Gabriel Ozorowski, Dennis R. Burton, Beatrice H. Hahn, George M. Shaw, Andrew B. Ward, Max Crispin, Christopher O. Barnes, Raiees Andrabi

## Abstract

Stabilizing the HIV envelope (Env) trimer in its native prefusion conformation is key to eliciting broadly neutralizing antibodies (bnAbs). We present a generalizable single-chain trimer-(SCT) design platform that enables the production of stable, native-like Env trimers across diverse HIV strains. Using a structure-guided approach, we developed CAP256.SU-SCT9, based on a CAP256.SU Env shown to induce V2-apex bnAbs in humans and rhesus macaques. CAP256.SU-SCT9 trimer exhibited native-like structure, bnAb-specific antigenicity, authentic glycosylation, and compatibility with nucleic acid and nanoparticle delivery. We extended this design strategy to a panel of HIV Envs representing global diversity, and immunogens known to initiate bnAb responses in humans. Structural analyses confirmed that SCTs consistently maintained prefusion-closed conformations, favorable glycan profiles, strong binding to trimer-specific bnAbs, and low reactivity with non-neutralizing antibodies. These findings establish SCT as a versatile platform for HIV trimer immunogen design, with potential to support next-generation vaccines aimed at eliciting protective bnAb responses.

## Introduction

The design of native-like HIV envelope (Env) spikes in a prefusion configuration is crucial for vaccine development, as it preserves the structural integrity and antigenic properties of the HIV Env glycoprotein (He et al., 2018; Joyce et al., 2017; Kong et al., 2016). Stabilizing Env ensures that it mimics the conformation of the spike on the virion surface, which is preferentially recognized by broadly neutralizing antibodies (bnAbs) (McCoy and Burton, 2017). HIV Env is highly metastable and prone to adopting non-native open conformations, exposing irrelevant or immunodominant epitopes, which are undesirable for effective immune responses (Burton and Hangartner, 2016; Havenar-Daughton et al., 2017). Stabilizing Env in its native-like, prefusion trimeric form allows for more effective presentation of key neutralizing epitopes, enhancing B cell recognition and directing immune responses toward broadly neutralizing antibody (bnAb) induction. This design approach increases the likelihood of generating protective, broadly reactive antibody responses – an important and critical goal in HIV vaccine development.

The HIV Env spike is a heterotrimer composed of gp120 and gp41 subunits that are derived from a gp160 precursor and are non-covalently linked (Harrison, 2015; Wyatt and Sodroski, 1998). To stabilize Env in a native-like prefusion trimeric conformation, several trimer stabilization strategies have been developed, including stabilized, soluble Env trimeric ectodomains (SOSIP - protein stabilized by SOS - disulfide bond, and IP - I559P mutation) (Sanders et al., 2013), NFL (native flexibly linked) (Sharma et al., 2015), UFO (uncleaved prefusion-optimized) (He *et al*., 2018), and RnS (repair-and-stabilization) (Rutten et al., 2018) designs. The SOSIP strategy stabilizes the HIV Env trimer by covalently linking the gp120 and gp41 subunits and replacing their native cleavage site with a hexa-arginine furin cleavage sequence. Additionally, a proline mutation in gp41 is introduced to enhance stability in the prefusion conformation(Sanders *et al*., 2013). This design has produced well-ordered trimers that elicit autologous neutralizing responses in preclinical models (Sanders and Moore, 2017; Sanders et al., 2015). Additional SOSIP modifications have sought to increase stability (Steichen et al., 2016), reduce immunodominant off-target B cell responses (de Taeye et al., 2015; Kulp et al., 2017), and minimize CD4-induced conformational changes (Chuang et al., 2017). However, SOSIP trimers remain prone to degradation, likely due to the furin cleavage site between gp120 and gp41 and the exposure of the trimer base (Turner et al., 2021). The NFL design strategy replaces the native furin cleavage site and introduces a flexible linker between gp120 and gp41, enhancing trimer integrity and stability (Guenaga et al., 2017; Sharma *et al*., 2015). The UFO design strategy replaces the metastable HR1 region of gp41 with a computationally designed shortened loop to improve trimer stability and antigenicity (He *et al*., 2018). Lastly, the RnS design strategy incorporates multiple mutations throughout the Env trimer to further reduce premature transition to an open conformation, improve folding and expression and enhance thermostability (Rawi et al., 2020; Rutten *et al*., 2018). Each of these strategies have advanced Env-based immunogen design, but challenges remain in consistently eliciting bnAb responses, requiring further refinements to optimize stability, immunogenicity, and epitope presentation.

There are two major strategies for designing vaccines to elicit bnAbs against HIV: the Env– antibody (Env–Ab) lineage-based approach (Haynes et al., 2012; Haynes et al., 2016), and the germline-targeting approach (Andrabi et al., 2018; Jardine et al., 2013; McGuire et al., 2013; Stamatatos et al., 2017; Steichen *et al*., 2016). The Env–Ab lineage-based approach begins with a defined unmutated common ancestor (UCA) of a known bnAb B cell lineage (Bhiman et al., 2015; Doria-Rose et al., 2014; Landais et al., 2017; Liao et al., 2013). A priming Env immunogen that binds the UCA is used to initiate the response, followed by a sequence of immunogens designed to guide antibody maturation by mimicking the natural co-evolution of Env and antibody. In contrast, the germline-targeting approach starts with a priming immunogen engineered to engage a broad set of naïve B cell precursors sharing key germline features. Boost immunogens are then sequentially administered to selectively expand and mature these clones toward the desired bnAb response. Both approaches utilize native-like trimer immunogens for boosting and/or prime-boosting within a multi-stage, multi-component HIV vaccine strategy (Andrabi *et al*., 2018; Burton and Hangartner, 2016; Caniels et al., 2025; Kwong et al., 2020; Steichen et al., 2019). To expand the repertoire of well-folded HIV boosting immunogens that capture global Env diversity and are compatible with various vaccine platforms, we developed here a new single-chain trimer (SCT) design strategy. This design builds upon previous trimer stabilization approaches by integrating multiple strategies—each of which individually yielded suboptimal results—and incorporating additional structure-guided modifications to enhance prefusion stability. We demonstrate that this SCT platform is generalizable across a diverse array of HIV-1 Envs, supporting its utility for vaccine development. As a proof of concept, we first focused on CAP256.SU Env, an Env-Ab lineage-based immunogen that has been shown to elicit V2-apex bnAbs in both human natural infection and rhesus macaque SHIV model (Bhiman *et al*., 2015; Doria-Rose *et al*., 2014; Roark et al., 2025; Roark et al., 2021). Through iterative structure-guided design, we developed CAP256.SU-based SCT trimers with enhanced prefusion stability, yielding improved antigenic and biophysical properties compared to previous CAP256.SU design (Gorman et al., 2020). Cryo-EM analysis confirmed that the resulting SCT trimers adopt a native-like prefusion structure. To assess generalizability, we applied the SCT design to a panel of HIV-1 Envs representing global group M diversity (deCamp et al., 2014; Hraber et al., 2018), as well as to Env–Ab lineage-based trimers that target distinct bnAb epitopes and have been shown to induce bnAbs in humans. These SCT immunogens consistently maintained native-like properties, as confirmed by antigenic profiling with bnAbs and non-neutralizing antibodies, negative-stain EM, and site-specific glycan analysis. Moreover, they are compatible with delivery as nucleic acid vaccines (mRNA, self-amplifying RNA (saRNA), DNA) or as protein-based nanoparticles, with minimal alteration to antigenic profiles. Together, these results establish the SCT strategy as a versatile and broadly applicable platform to support iterative HIV vaccine design and bnAb discovery through reverse vaccinology (Burton, 2017).

## Results

### Design of native-like single chain trimers for HIV CAP256 Env based immunogens

The V2-apex epitope on the HIV envelope is a promising target for vaccine design, as bnAbs directed at this site tend to be both potent and broad. These bnAbs are commonly elicited in individuals who develop bnAb responses during natural infection and typically require lower levels of somatic hypermutation compared to bnAbs targeting other epitopes (Andrabi et al., 2015; Landais et al., 2016). The CAP256.SU Env, or its derivative CAP256.wk34, has been shown to initiate the CAP256.01–33 bnAb lineage during natural infection in a human (Bhiman *et al*., 2015; Doria-Rose *et al*., 2014). Similarly, rhesus macaques infected with CAP256.SU SHIV frequently develop V2-apex bnAb responses (Habib R, 2025; Roark *et al*., 2025; Roark *et al*., 2021). Based on this evidence, we aimed to design a B cell lineage-based immunogen derived from CAP256.SU Env to induce V2-apex bnAbs through vaccination in preclinical models. However, existing trimer stabilization strategies—such as DS-SOSIP, NFL, and RnS—failed to consistently produce well-folded CAP256.SU trimers. This limitation prompted us to develop a new Env trimer design strategy that integrates key stabilizing mutations from multiple existing approaches, with the goal of producing a design adaptable to CAP256.SU and generalizable across diverse HIV Env strains.

To generate a prefusion-stabilized CAP256.SU Env trimer, we employed a combinatorial mutational strategy incorporating elements from the DS-SOSIP, NFL-TD8, and RnS designs (Chuang *et al*., 2017; Guenaga et al., 2015; Rutten *et al*., 2018). Our goal was to optimize key trimer subdomains to enhance hydrophilic interactions at the apex, limit CD4-induced conformational changes, and reduce exposure of the non-neutralizing V3 loop face. We introduced stabilizing mutations in both the gp120 subunit (including V1V2 and V3 regions) and the gp41 subunit (Figure 1A–B). Overall, we aimed to develop a trimer platform that would be resistant to cleavage-dependent degradation and compatible with nucleic acid and self-assembling nanoparticle delivery systems. To achieve this, we systematically combined stabilized subdomain variants and covalently linked gp120 and gp41 using a flexible glycine-serine linker. Trimer formation was assessed by size-exclusion chromatography (SEC). The final construct—a covalently linked gp120-gp41 trimer—is referred to as a Single-Chain Trimer (SCT).

**Figure 1.**
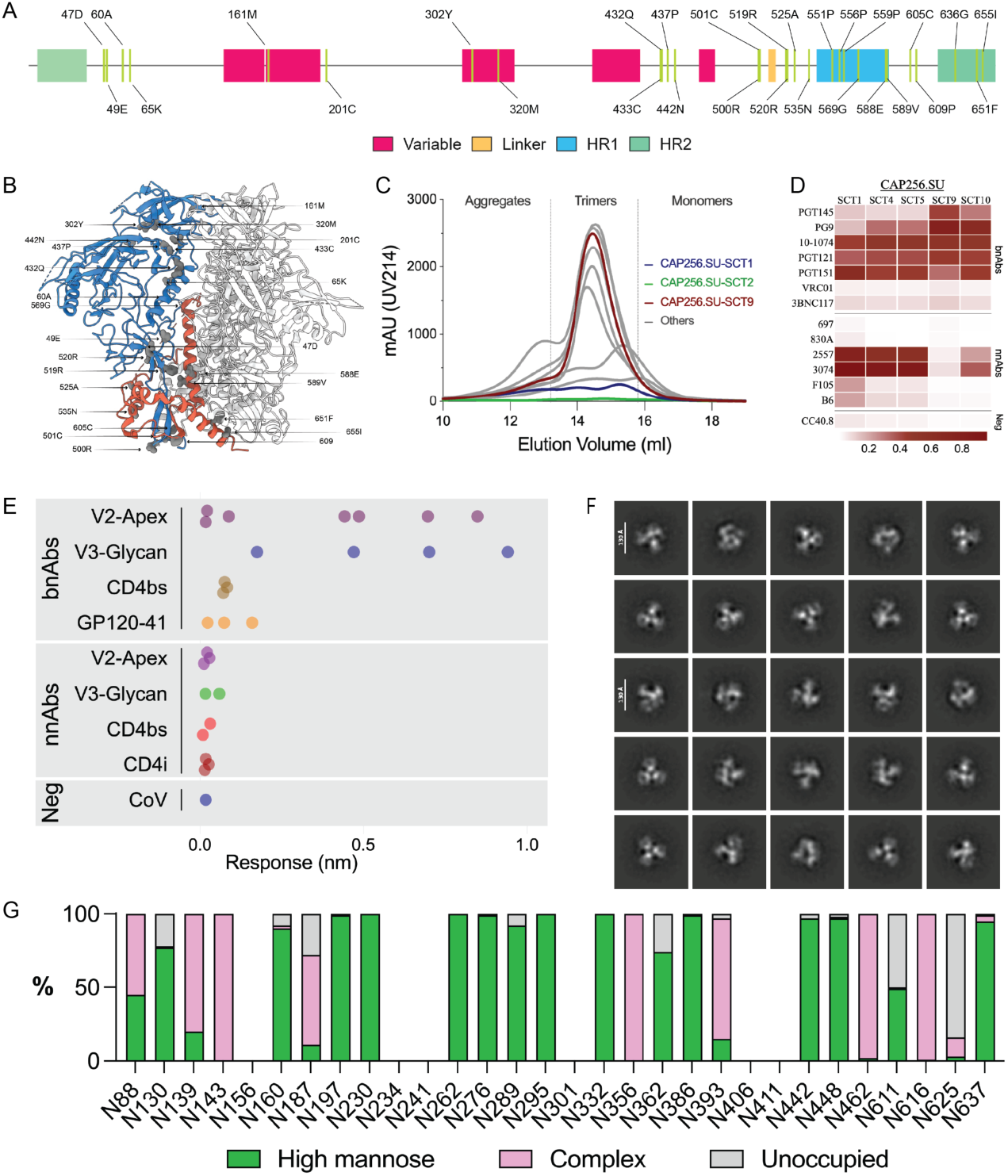
Design and characterization of a single-chain trimer (SCT) based on CAP256.SU Env to enhance expression of native-like soluble trimers. **A.** Gene map illustrating point mutations incorporated into the SCT design. SCT-specific mutations are shown in grey. Heptad repeat regions HR1 and HR2 are highlighted in blue and green, respectively, alongside other distinct regions within the CAP256.SU Env. **B.** Schematic representation of SCT mutations mapped onto the CAP256.SU Env. The gp120 region is highlighted in blue, gp41 in red, and SCT mutations are indicated in grey. **C.** Size exclusion chromatography (SEC) profiles of CAP256.SU-SCT1 to SCT10 on a Superdex 200 column. The base construct CAP256.SU-DS-SOSIP (SCT1) is shown in blue, CAP256.SU-NFL (SCT2) in green, and CAP256.SU-SCT9 in red. Dashed lines indicate separation of trimeric forms from aggregated or dimeric/monomeric Env species. **D.** Heatmap showing the maximum BioLayer Interferometry (BLI) binding response of selected well-assembled CAP256.SU-SCTs (100 nM) to a panel of HIV-1 bnAbs and non-neutralizing antibodies (nnAbs). CC40.8, a SARS-CoV-2 stem-helix antibody, is included as a negative control. **E.** Dot plot displaying the maximum BLI binding response of CAP256.SU-SCT9 (100 nM) to an expanded panel of HIV-1 bnAbs and nnAbs. **F.** Negative-stain electron microscopy (nsEM) 2D class averages of CAP256.SU-SCT9 reveal that the trimers adopt a 100% native-like conformation. **G.** Glycan occupancy profile assessed by site-specific glycan analysis (SSGA) using proteomics. High-mannose glycans are shown in green, complex glycans in pink, and unoccupied sites in grey.

The initial construct, CAP256.SU-SCT1, incorporated the SOSIP mutations (501C–605C and I559P), an RnS mutation at position 60A, and a disulfide bond between residues 201C and 433C to minimize CD4-induced epitope exposure. This core set of mutations was retained across all ten design variants (CAP256.SU-SCT1 through CAP256.SU-SCT10). Subsequent iterations introduced additional stabilizing mutations in both gp41 and gp120. For example, CAP256.SU-SCT2 (SCT2) incorporated mutations derived from the NFL-TD8 design. These included two gp120 mutations—432Q and 500R—as well as several gp41 mutations: glycine substitutions at positions 569 and 636 to reduce the likelihood of extended helices forming at the helix-to-coil transitions in HR1 and HR2; a Q551P mutation within the metastable HR1 loop to limit coil-to-post-fusion helix triggering; and a double mutation, 519R-520R, aimed at decreasing the hydrophobicity of the the solvent exposed segment of the fusion peptide. CAP256.SU-SCT3 (SCT3) included two RnS-derived gp120 mutations (437P and 442N) and nine RnS-derived gp41 mutations (525A, 535N, 556P, 588E, 589V, 609P, 651F, 655I, and 658V). CAP256.SU-SCT4 (SCT4) combined all four gp120 and fourteen gp41-stabilizing mutations from SCT2 and SCT3. CAP256.SU-SCT5 (SCT5) retained the full mutation set from SCT4 but replaced the 501C–605C disulfide bond with a 501C–663C linkage to further stabilize the trimer base. To enhance gp120 stability, six additional NFL-TD8-derived mutations (47D, 49E, 65K, 161M, 302Y, and 320M) were introduced. These were systematically incorporated into the following constructs: CAP256.SU-SCT6 = SCT1 + gp120 stabilizing mutations, CAP256.SU-SCT7 = SCT2 + gp120 stabilizing mutations, CAP256.SU-SCT8 = SCT3 + gp120 stabilizing mutations, CAP256.SU-SCT9 = SCT4 + gp120 stabilizing mutations, CAP256.SU-SCT10 = SCT5 + gp120 stabilizing mutations (Figure S1A). This iterative design strategy progressively enhanced both the structural stability and antigenic properties of the CAP256.SU Env trimer constructs.

To evaluate improvements in prefusion trimer assembly and folding of soluble Env trimers, each construct was expressed in mammalian 293F cells and purified using *Galanthus nivalis* lectin (GNL) affinity chromatography, followed by SEC. GNL binds to Env glycans and enables an unbiased purification of all Env species—including trimers, dimers, monomers, aggregates, and truncated forms—thereby avoiding potential selection bias introduced by antibody-based affinity purification. SEC profiles revealed that SCT4, SCT5, SCT9, and SCT10 exhibited markedly improved yields (Figure S1B) and a predominance of well-formed trimers, with minimal aggregation and unfolded monomers, in contrast to CAP256.SU-SOSIP (SCT1) and CAP256.SU-NFL (SCT2) (Figure 1C). To identify the construct that most closely resembled the native prefusion conformation of CAP256.SU Env, we assessed antigenic profiles using biolayer interferometry (BLI) with a panel of bnAbs and non-neutralizing antibodies (nnAbs) targeting diverse HIV Env epitopes. Among all designs, CAP256.SU-SCT9 demonstrated the strongest binding to PG9 and PGT145, V2-apex bnAbs that selectively recognize the quaternary epitope formed on closed native-like trimers (Figure 1D). Additionally, CAP256.SU-SCT9 exhibited robust binding to bnAbs targeting the V3 glycan site (PGT121, 10-1074), the CD4 binding site (VRC01, 3BNC117), and the gp120–gp41 interface (PGT151, VRC34.01), while showing minimal binding to nnAbs targeting various Env surfaces (Figure 1D).

To further assess the antigenic properties of CAP256.SU-SCT9, we conducted a BLI binding assay using an expanded panel of bnAbs and nnAbs targeting diverse HIV Env epitopes. The results confirmed that CAP256.SU-SCT9 is a well-assembled trimer, showing enhanced binding to bnAbs across multiple epitope classes while eliminating binding to nnAbs—consistent with a closed, native-like trimer conformation (Figure 1E). nsEM 2D classification analysis further validated that CAP256.SU-SCT9 consistently adopts the native prefusion trimer architecture (Figure 1F). In addition, mass spectrometry-based glycan profiling revealed a predominance of high-mannose and complex glycans, with minimal presence of non-native glycan structures (Figure 1G), supporting proper glycan processing and structural integrity. Based on its superior biophysical and antigenic characteristics, CAP256.SU-SCT9 was selected as the lead trimer construct that most closely mimics the native prefusion conformation of Env. This CAP256.SU-SCT9 design also served as the foundation for a follow-up study involving a CAP256.SU Env lineage-based, germline-targeting trimer. Using CAP256.SU-SCT9 as the base construct, the engineered trimer immunogen—named CAP256.SU-SCT9-GT1, or CAP256-GT1— demonstrated substantially improved binding to multiple diverse V2-apex bnAb precursors. CAP256-GT1 effectively primed rare V2-apex long CDRH3 precursor B cells and generated a robust memory response, which ultimately elicited potent bnAb responses in outbred rhesus macaques (Li X et al., 2025, in preparation).

Using this same iterative design strategy, we engineered ten additional constructs (SCT11– SCT20) based on the CAP256.WK34 Env sequence (Figures S1C-D, S2A-B). Characterization by SEC and antigenicity profiling identified CAP256.wk34-SCT19 as the most promising candidate, harboring the same combination of mutations as CAP256.SU-SCT9 (Figure S2C-E). This result supports the broader applicability of this design strategy to potentially stabilize diverse HIV Env strains in a prefusion-closed state.

### CAP256.SU-SCT9 trimer is suitable for nucleic acid-based vaccine delivery

To evaluate the suitability of the engineered native-like CAP256.SU-SCT9 trimer for nucleic acid-based vaccine delivery, such as mRNA lipid nanoparticles (LNPs), self-amplifying mRNAs and DNA, we assessed its antigenic profile when expressed as a membrane-bound trimer on mammalian cells using a cell surface binding assay (CELISA). A comprehensive panel of HIV Env mAbs was used, including 21 human and rhesus V2-apex bnAbs, 6 V3-glycan bnAbs, 9 CD4bs bnAbs, and 4 gp120–gp41 interface bnAbs. The panel also included 15 nnAbs, comprising 4 V2-apex nnAbs, 5 V3 nnAbs, 3 CD4bs nnAbs, and 3 CD4i nnAbs (Figure. 2A, S3). CELISA binding data revealed a favorable antigenic profile consistent with a native-like membrane-bound CAP256.SU-SCT9 trimer. The membrane-bound CAP256.SU-SCT9 construct was strongly recognized by trimer-dependent V2-apex bnAbs, as well as bnAbs targeting the V3-glycan and CD4bs epitopes, while exhibiting little to no binding to nnAbs targeting various Env regions—except for some linear V3-directed nnAbs (Figure 2A, S3). These findings were consistent with BLI binding profiles of the corresponding soluble trimer (Figure 2B, S3) and with pseudovirus neutralization IC₅₀ values (Figure 2C, S3). A slight discrepancy was observed between the cell surface-expressed and soluble trimers in binding to linear V3 loop-targeting nnAbs. We attribute this to a minor subpopulation of non-native open trimer forms on the cell surface. Importantly, binding data from the membrane-bound SCT9 trimer strongly correlated with both BLI measurements of the soluble trimer (Fig. 2D) and IC_50_ neutralization titers against the CAP256.SU pseudovirus (Fig. 2F). Furthermore, BLI binding of the soluble trimer also correlated well with pseudovirus neutralization IC₅₀ values to pseudovirus lacking stabilization mutations (Figure 2E), reinforcing the structural and functional integrity of CAP256.SU-SCT9 in both soluble and membrane-anchored formats. Together, these results validate the CAP256.SU-SCT9 design as suitable for both nucleic acid-based and protein-based vaccine delivery platforms.

**Figure 2.**
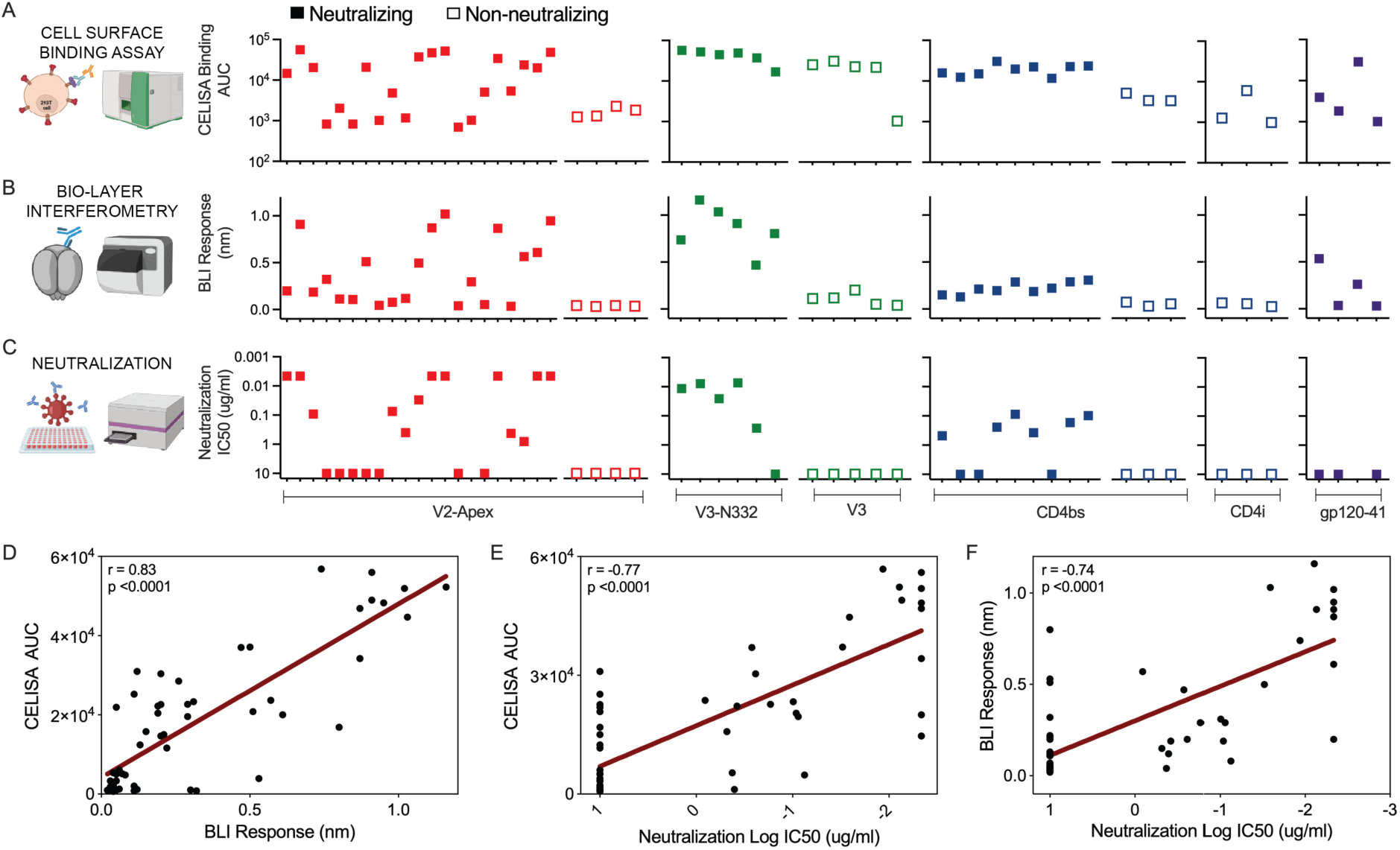
Antigenic profile of soluble and membrane-bound CAP256.SU-SCT9 trimers. **A.** Dot plots showing cell surface-expressed CAP256.SU-SCT9 trimer binding (CELISA), represented as Area Under the Curve (AUC) values, to a panel of HIV bnAbs and non-neutralizing antibodies (nnAbs) targeting diverse Env epitopes. **B.** Dot plots showing BioLayer Interferometry (BLI) maximum binding responses of the soluble CAP256.SU-SCT9 trimer (at 500 nM) to the same panel of HIV bnAbs and nnAbs targeting various Env epitopes. **C.** Dot plots presenting reciprocal neutralization IC_50_ titers of the HIV bnAbs and nnAbs against a CAP256.SU Env-pseudotyped virus, demonstrating antibody potency across different Env targets. **D.** Correlation analysis between the cell surface binding assay AUC values and the BLI maximum binding responses. A strong positive correlation was observed (correlation coefficient, r = 0.83, p < 0.0001), indicating that soluble trimer binding closely mirrors membrane-bound trimer recognition. **E.** Correlation between cell surface binding AUC values and neutralization potency, expressed as log IC_50_. A significant negative correlation (r = -0.77, p < 0.0001) indicates that higher cell surface binding corresponds to more potent neutralization. **F.** Correlation between BLI maximum binding responses and neutralization potency (log IC_50_ values). BLI binding also strongly correlates with neutralization activity (r = -0.74, p < 0.0001), supporting the relevance of soluble trimer binding measurements to functional antibody responses.

### CAP256.SU-SCT9 trimer is compatible with self-assembling nanoparticles that exhibits enhanced antigenicity

To assess the ability of the CAP256.SU-SCT9 native-like trimer to form a self-assembling multivalent nanoparticle with enhanced antigenicity, we grafted the trimer onto a self-assembling ferritin scaffold. The ferritin scaffold, derived from *Helicobacter pylori* (Kanekiyo et al., 2013; Sliepen et al., 2015), naturally forms a 24-subunit nanoparticle and thus displays eight copies of the CAP256.SU-SCT9 trimer (Figure. 3A). These trimer-ferritin nanoparticles were expressed in the 293F mammalian expression system and purified using PGT145 antibody affinity chromatography. Size-exclusion chromatography of the purified proteins showed a single major peak at an elution volume of ∼10 mL on a Superose 6 column, consistent with the expected size of the self-assembling trimer-ferritin nanoparticles (Figure 3B). To evaluate the antigenicity of the CAP256.SU-SCT9 trimer nanoparticle, we performed BLI binding assays using a panel of HIV Env-specific bnAbs and nnAbs targeting various epitopes. The trimer nanoparticle showed markedly enhanced binding to bnAbs compared to the soluble trimer, likely due to avidity effects from multivalent display (Figure 3C-D). Notably, enhanced binding was observed across multiple bnAb epitopes without a corresponding increase in binding by nnAbs, indicating selective improvement in bnAb epitope antigenic presentation.

**Figure 3.**
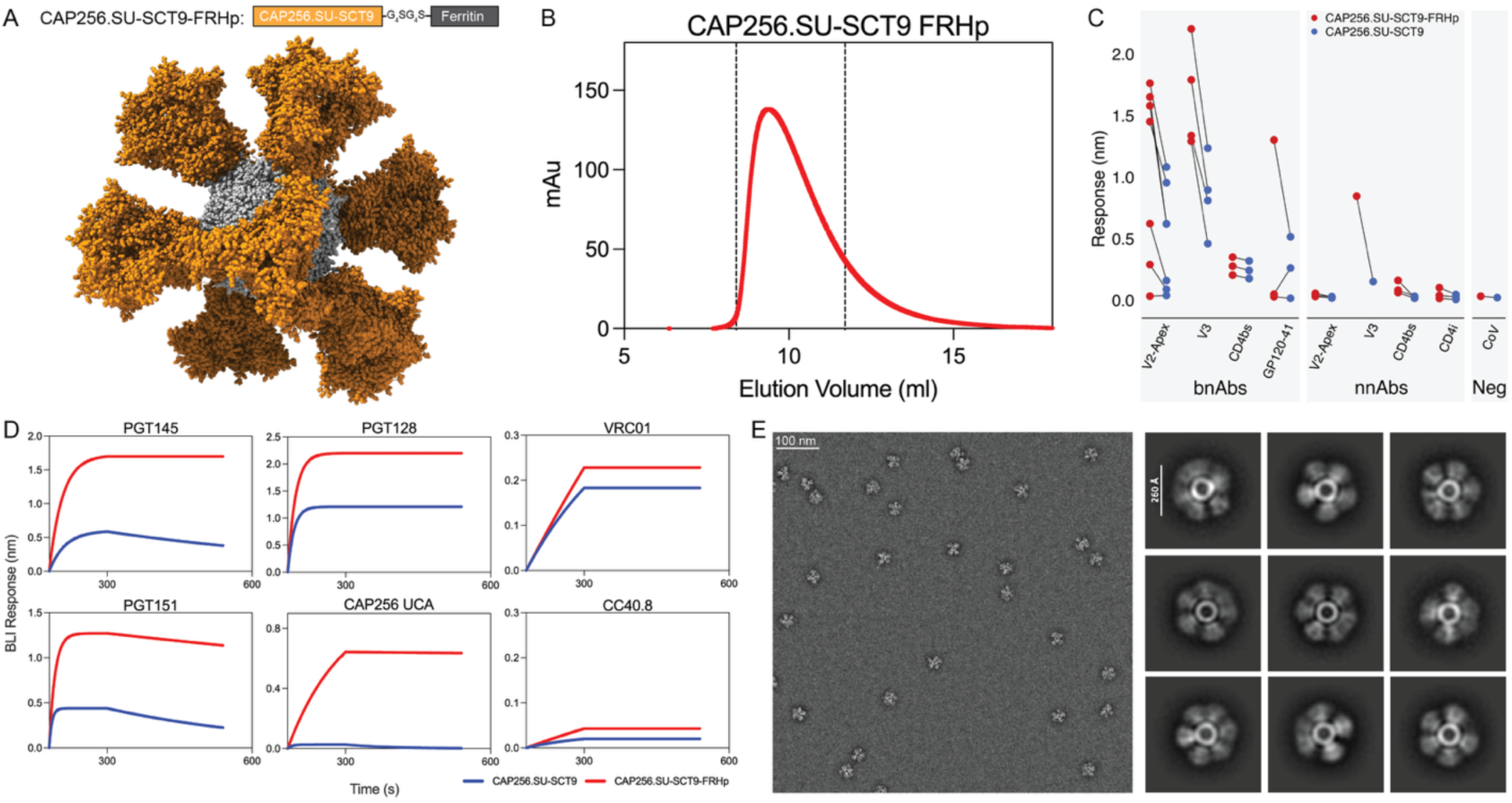
Design and characterization of self-assembled ferritin nanoparticles displaying CAP256.SU-SCT9 trimers for enhanced antigenicity. **A.** Schematic representation of the CAP256.SU-SCT9-FRHp ferritin nanoparticle design. Each nanoparticle displays 8 copies of the CAP256.SU-SCT9 trimer (highlighted in green) arranged on the surface of a 24-subunit ferritin core (shown in grey), allowing multivalent presentation of the Env trimer. **B.** Size exclusion chromatography (SEC) profile of the CAP256.SU-SCT9-FRHp nanoparticle, following purification by PGT145 affinity chromatography and subsequent SEC on a Superose6 column. A single nanoparticle peak eluting at ∼10 mL is shown within the dashed lines, indicating a uniform and stable nanoparticle population. **C.** Dot plot comparing the maximum BLI binding responses of the CAP256.SU-SCT9 soluble trimer (500 nM) and the CAP256.SU-SCT9-FRHp nanoparticle (500 nM) to a panel of HIV bnAbs and non-neutralizing antibodies (nnAbs) targeting diverse Env epitopes. Lines connect binding values for each antibody to both the trimer and the nanoparticle, highlighting differences in binding avidity. **D.** BLI binding kinetics (association and dissociation curves) comparing CAP256.SU-SCT9 trimer (500 nM, blue) and CAP256.SU-SCT9-FRHp nanoparticle (500 nM, red) to a representative set of HIV bnAbs, including V2-apex-targeting unmutated common ancestor (UCA) antibody, CAP256 UCA. CC40.8, a SARS-CoV-2 S2 stem-targeting antibody, serves as a negative control. The multivalent nanoparticle shows enhanced avidity binding and slower off-rates compared to the soluble trimer. **E.** Negative-stain electron microscopy (nsEM) of CAP256.SU-SCT9-FRHp nanoparticle. A representative micrograph (left) and corresponding 2D class averages (right) reveal well-formed, fully assembled 24-mer ferritin nanoparticles displaying uniformly distributed Env trimers.

We also tested the binding of the CAP256.UCA precursor (Doria-Rose *et al*., 2014)—a member of the CAP256.VRC26 V2-apex bnAb lineage—to the trimer nanoparticle (Figure 3D). Compared to the soluble trimer, the nanoparticle showed strong binding with a stable off-rate, suggesting high-avidity interaction with the unmutated ancestor version of the bnAb. Previous studies have revealed that this stable binding is associated with efficient rare B cell priming (Abbott et al., 2018; Dosenovic et al., 2018), supporting the potential of the CAP256.SU-SCT9 trimer nanoparticle as a candidate immunogen to initiate V2-apex bnAb lineages within an Env lineage-based vaccine approach.

To assess the structural integrity and homogeneity of the nanoparticle, we conducted nsEM analysis on PGT145-purified trimer nanoparticles. nsEM imaging revealed uniformly sized, well-folded particles with consistent morphology, confirming successful multivalent display and structural uniformity (Figure 3E). We additionally applied the same nanoparticle design strategy and assessments on CAP256.Wk34-SCT19, which produced similar results (Figure S4A-E). Altogether, these findings demonstrate that the SCT design enables the CAP256.SU-SCT9 and CAP256.Wk34-SCT19 constructs to assemble uniformly on a nanoparticle platform, conferring favorable antigenic and structural properties.

### Cryo EM Structure of CAP256.SU-SCT9 trimer

To evaluate whether the introduction of SCT9 stabilizing mutations maintains a native-like Env conformation, we obtained a 3.5 Å resolution cryo-EM structure of trimeric CAP256SU.SCT9 in complex with the antigen-binding fragment (Fab) of 40591-a.05 (Roark *et al*., 2025), a V2-apex-targeting neutralizing antibody derived from CAP256SU SHIV-infected rhesus macaques (Figure 4A,B, Figure S5) (Habib R, 2025; Roark *et al*., 2025; Roark *et al*., 2021) . At this resolution, we were able to model the backbone and sidechain positions for most residues of gp140. To more accurately model density at the Fab-Env interface, we also determined a 1.6 Å resolution crystal structure of the 40591-a.05 Fab that was docked into the corresponding density at the trimer apex. Additional stabilization of the SCT constructs was achieved through the introduction of gp120-stabilizing mutations 302Y and 320M, derived from the NFL-TD8 design (Guenaga *et al*., 2015). These mutations improved hydrophobic packing with gp120 residue 177Y and introduced a hydrogen bond between 302Y and 320M, thereby preventing V3 loop exposure and CD4-induced conformational change (Figure 4B). Superposition of the gp140 backbones from the CAP256SU.SCT9-40591-a.05 Fab structure with previously resolved structures of CAP256SU SOSIP (PDB: 6VRW) and the single-chain BG505 IDL NFL TD CC^+^ trimer (PDB: 9BEW) yielded a C_⍺_ root-mean-square deviations (RMSD) of 0.80 Å and 1.02 Å, respectively (Figure 4C) (Gorman *et al*., 2020; Rutten *et al*., 2018). Comparisons with Env structures derived from other stabilization strategies, such as RnS-SOSIPs, triple tandem trimers, and interdomain lock Env trimers (IDLs) also yielded low RMSDs (Figure S6) (del Moral-Sánchez et al., 2024; Rutten *et al*., 2018; Zhang et al., 2024). The overall global similarity between CAP256.SU-SCT9 and SOSIP-stabilized or single-chain Env trimers suggests that the mutations introduced during SCT design are sufficient to stabilize the prefusion, closed trimer and recapitulate native-like Env architecture.

**Figure 4.**
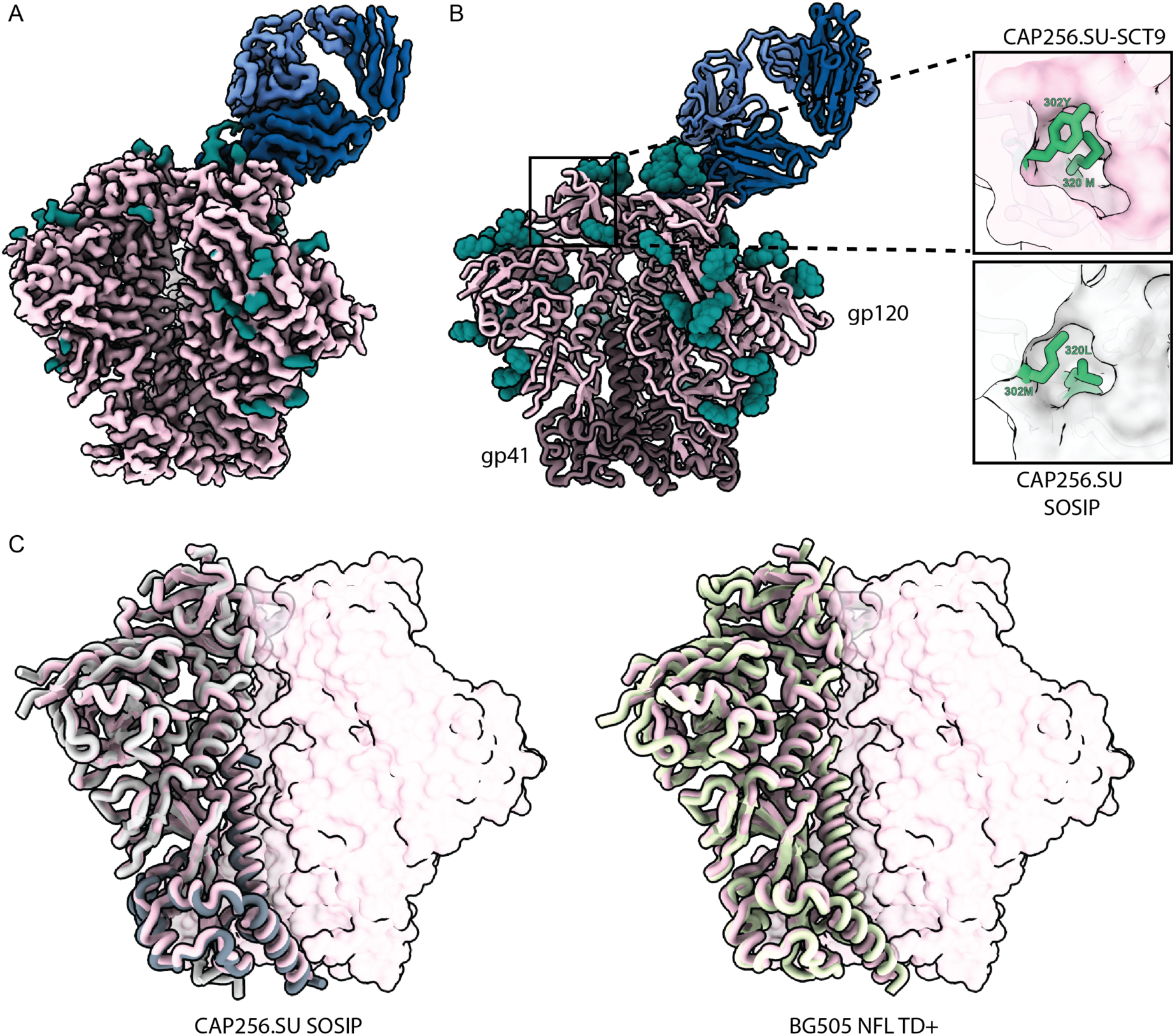
Cryo-EM volume map and structure of CAP256.SU-SCT9 in complex with rhesus-macaque derived V2-apex targeting bnAb Fab. **A, B.** Cryo-EM density resolved to 3.5A **(A)** and overall structure **(B)** of CAP256.SU-SCT9 (gp120 shown in light pink and gp41 shown in dark pink), glycans (shown in teal), in complex with rhesus-macaque derived V2-apex targeting bnAb Fab 40591-a.05 (shown in blue). Insets reveal that the additional NFL-TD8 derived gp120-stabilizing mutations introduced in later iterations of the SCT constructs (CAP256.SU-SCT6-10) **C.** Superposition of CAP256.SU-SCT9 with CAP256.SU SOSIP (shown in grey; PDB 6VRW) and BG505 NFL TD+ (shown in sage; PDB 6P62). RMSD (0.8 Å and 1.02 Å) calculations were determined from alignment of 402 and 393 C_⍺_ atoms of gp140 subunit, respectively.

### The Single Chain Trimer stabilization design is broadly applicable across diverse global HIV-1 subtypes

The successful stabilization of the CAP256.SU-SCT9 trimer prompted us to extend the SCT design strategy to a broader panel of HIV-1 Envs representing global viral diversity. Using the same trimer stabilization approach applied to CAP256.SU-SCT9, we generated soluble SCTs from global panels of diverse representative HIV-1 Env strains (Figure 5A) (deCamp *et al*., 2014; Hraber *et al*., 2018). These SCTs were purified via GNL affinity chromatography followed by Superdex 200 size-exclusion chromatography, without additional positive or negative selection using antibodies and yielded between 3.7-15 mg per liter of 293F transfection. SEC profiles revealed that all 18 SCTs from the global Env panel exhibited a single dominant peak corresponding to a trimeric species, indicating consistent folding and efficient trimer assembly with minimal aggregates or unassembled monomers (Figure 5B).

**Figure 5.**
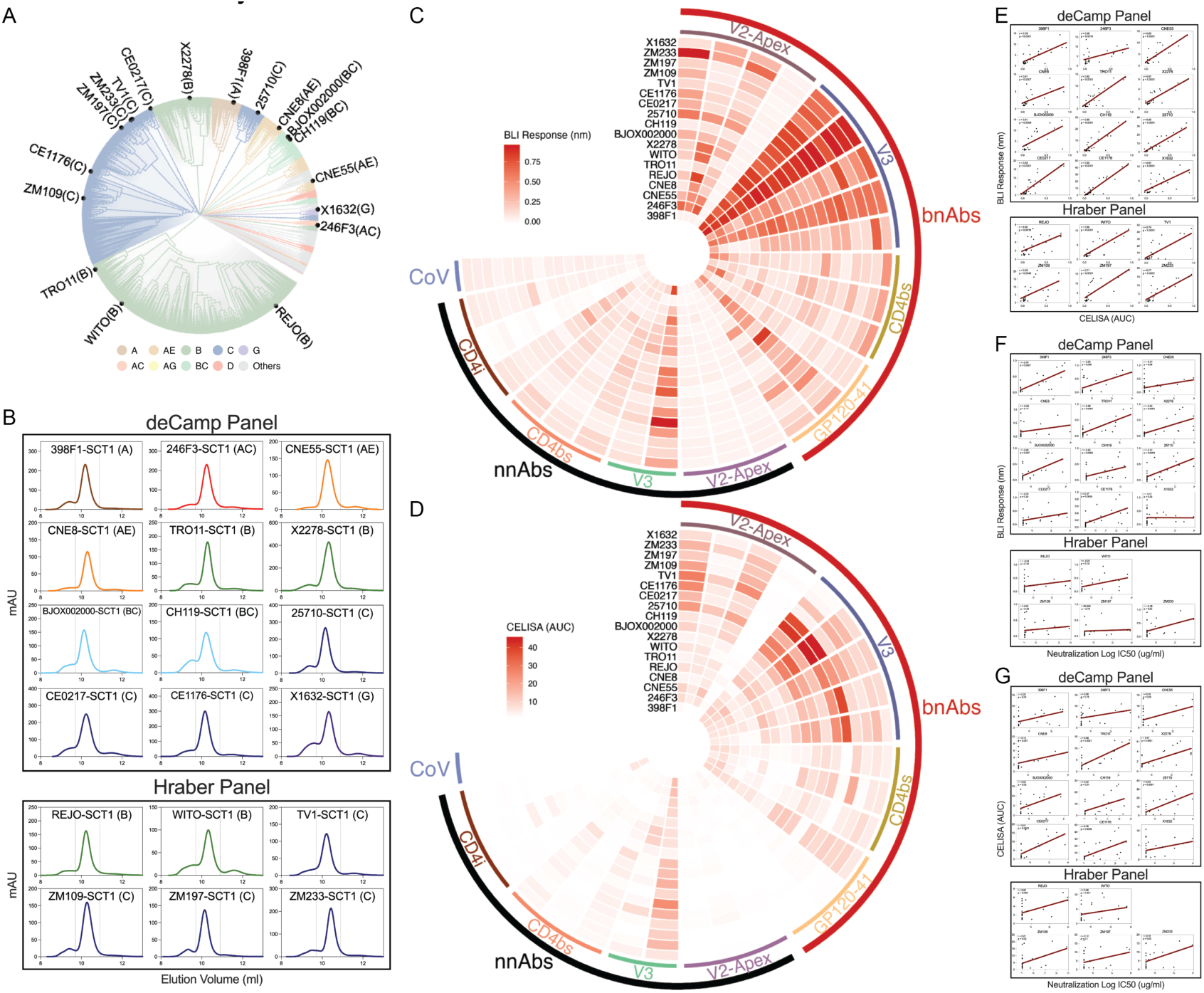
Design and characterization of SCTs representing global HIV-1 diversity. **A.** Phylogenetic tree illustrating the selected global panel of HIV-1 Env strains, representing a broad range of genetic diversity across major HIV-1 subtypes. This panel was chosen based on strains characterized in deCamp et al. and Hraber et al. to ensure coverage of global circulating strains for immunogen design. **B.** Size-exclusion chromatography (SEC) profiles of GNL-purified SCTs from the global panel, run on Superdex200 columns. A single major peak within the dashed region indicates consistently folded trimeric populations across the diverse SCT panel. **C.** Heatmap showing BLI maximum binding responses of the GNL-purified global SCT trimer panel (100 nM) to a panel of HIV-1 bnAbs and non-neutralizing antibodies (nnAbs) targeting diverse Env epitopes. Strong binding to bnAbs and minimal binding to nnAbs suggest favorable antigenic profiles and proper epitope presentation. **D.** Heatmap of cell surface binding responses (CELISA) measuring transmembrane expression of the global SCT panel in HEK293T mammalian cells. Binding to the same panel of bnAbs and nnAbs confirms consistent membrane-bound trimer expression and antigenicity across subtypes. **E.** Linear regression analysis (red line) showing a strong positive correlation between BLI maximum binding responses and CELISA AUC values, supporting concordance between soluble and membrane-bound trimer antigenicity. **F.** Linear regression showing a moderate negative correlation between BLI maximum binding responses and neutralization potency (log IC_50_), indicating that higher binding affinity corresponds to increased neutralization sensitivity. **G.** Linear regression showing a strong negative correlation between CELISA AUC values and log IC_50_, suggesting that stronger membrane-bound trimer binding is predictive of improved neutralization sensitivity.

We next evaluated the antigenicity of the global panel SCTs using BLI, cell surface binding assays, and virus neutralization assays against a focused panel of bnAbs and nnAbs targeting multiple Env epitopes. BLI profiling demonstrated that most SCTs bound strongly to trimer-dependent V2-apex and V3-glycan bnAbs and bnAbs to other Env site, while showing minimal binding to nnAbs (Figure 5C). To improve the trimer quality, we positively selected the global panel SCTs through PGT 145 affinity column chromatography. The yields varied from 300ug to 4 mg per liter of 293F transfection (Figure S7A). We applied the same method to profile the PGT145 purified global panel SCTs. The overall BLI binding profile was similar regardless of PGT145 selection, however, the positive selection led to enhanced affinity to V2-Apex and V3 bnAbs and further reduced binding to nnAbs (Figure S7B). These results indicate that the global panel SCTs predominantly adopt a native-like, prefusion-closed trimer conformation and confirm the broad adaptability of our SCT strategy across diverse HIV-1 strains. To further validate these findings, we assessed the antigenic profiles of the SCTs by CELISA, which corroborated well with the BLI results for the soluble trimers (Figure 5D). We then examined the correlation between BLI binding, CELISA responses, and neutralization IC₅₀ titers. For most constructs, strong to modest correlations were observed between antigen binding and neutralization potency, supporting the notion that these SCTs maintain native-like antigenicity consistent with functional virus-associated spikes (Figure 5E-G and Figure S7C). However, a few Envs - such as REJO and X1632 - showed weaker correlations, suggesting that in certain strain backgrounds, the SCT-stabilizing mutations may not fully recapitulate the native virion spike configuration. These differences may stem from conformational variability or altered glycan processing between the soluble SCTs and their corresponding viral spikes.

Finally, to confirm that SCT mutations consistently stabilize the diverse set of Envs in a native-like prefusion, closed state, we analyzed the global panel of SCTs, purified via PGT145 antibody affinity chromatography, using nsEM (Figure 6). Across clades, the majority of nsEM 2D class averages showed properly folded, trimeric Envs stabilized by the SCT design. Native Envs on the viral surface, as well as stabilized SOSIP constructs, inherently fluctuate between open-occluded and prefusion closed conformations (Ozorowski et al., 2017; Wang et al., 2018). While individual constructs vary in flexibility, as exemplified by the prominent apex hole observed in 25710 SCT and 246F3 SCT, which remains less noticeable in other constructs, the 2D class averages indicate that SCT mutations consistently shift the equilibrium of all Envs in the panel toward the prefusion closed state.

**Figure 6.**
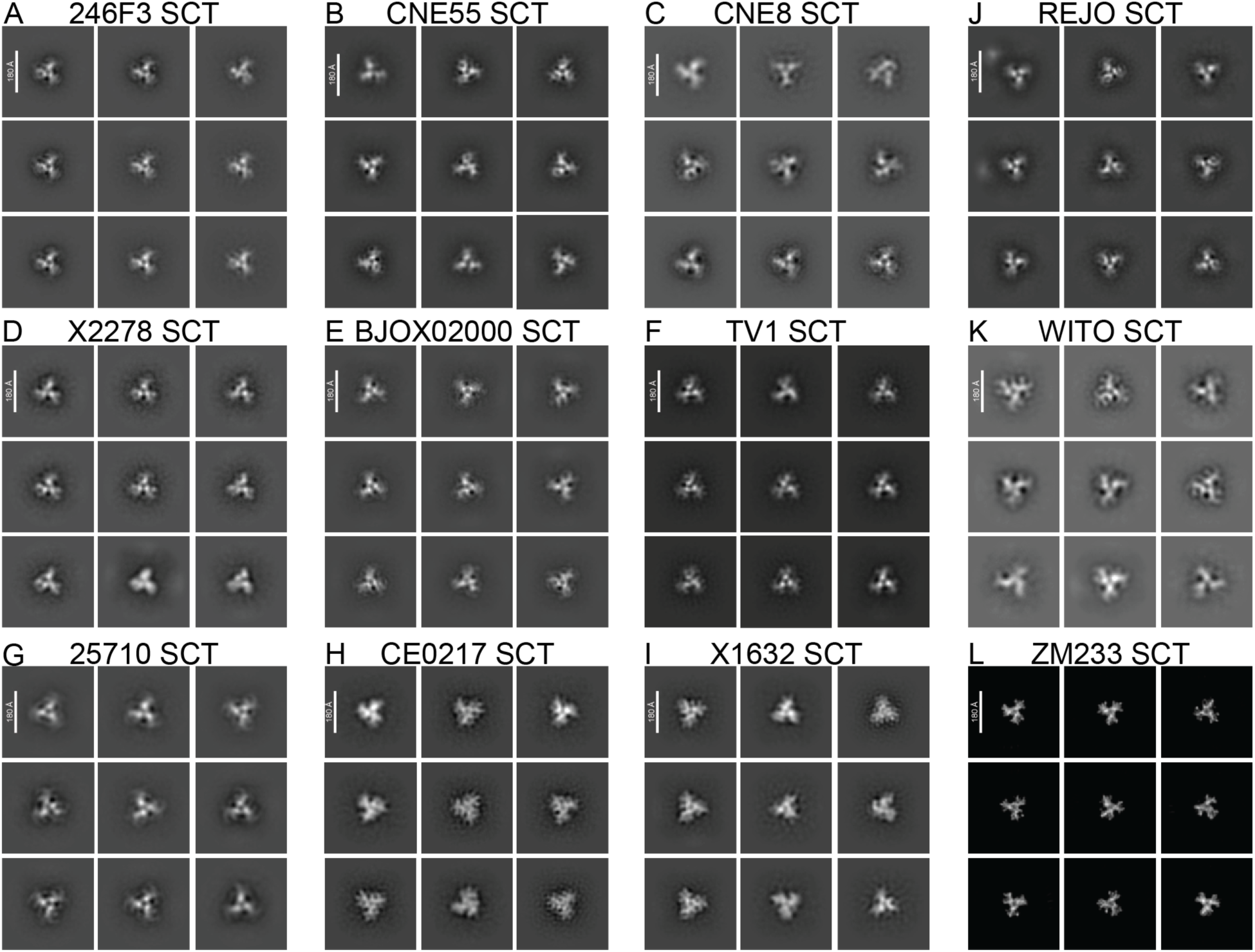
Negative-stain electron microscopy (nsEM) analysis of SCT-stabilized Env trimers representing global HIV-1 diversity. Representative 2D class averages are shown for selected single-chain trimer (SCT)-stabilized HIV-1 Env glycoproteins from the global panel. The images reveal that the majority of the Env trimers adopt closed or partially open native-like conformations, characteristic of well-folded, prefusion structures. These structural features are consistent across diverse HIV-1 subtypes, supporting the structural integrity of the SCT design across global panel. The scale bar represents 180 Å.

Overall, our findings demonstrate that the SCT design strategy is broadly generalizable beyond the CAP256.SU strain. This approach can be successfully applied to a diverse panel of HIV-1 Envs representing global subtype diversity. The consistent trimeric assembly and favorable antigenic profiles observed across this panel underscore the SCT platform’s robustness and utility for vaccine design. Moreover, the intrinsic structural stability of SCTs makes them well-suited for integration into advanced delivery systems, including mRNA-based vaccine platforms.

### Single Chain Trimers mimic native-like spike glycan profiles

N-linked glycans on HIV trimers play an important and essential role in proper protein folding, trimer stabilization, and antibody recognition. To determine if the designed SCT constructs resemble native-like HIV trimers in their glycoprotein composition glycan shield, we performed site-specific glycosylation and occupancy of the soluble global panel SCT proteins purified by PGT145 affinity chromatography (Behrens et al., 2016; Struwe et al., 2018). Glycosylation was determined by enzymatic digestion with serine proteases (trypsin, chymotrypsin and α-lytic protease) in three independent experiments and resulting peptides were analysed by liquid chromatography-mass spectrometry (LC-MS). Overall, an average of 62% of predominant oligomannose-type glycan occupancy was observed in all the global panel SCTs, without significant differences across a wide variety of HIV subtypes, ranging from 51.7% (BJOX002000 SCT) to 70% (WITO SCT) (Figure 7A). Further investigation into specific domains determined that the glycan occupancy on gp41 was reduced as expected, demonstrating an average of 33% of oligomannose composition and ranging from 56.7% (CNE55 SCT) to 14.7% (BJOX002000 SCT). This difference is likely largely due to very low or no oligomannose occupancy at sites N611, N616 and N625, while glycan site N637 is predominantly occupied by high mannose glycans (Figure 7C and Figure S8).

**Figure 7.**
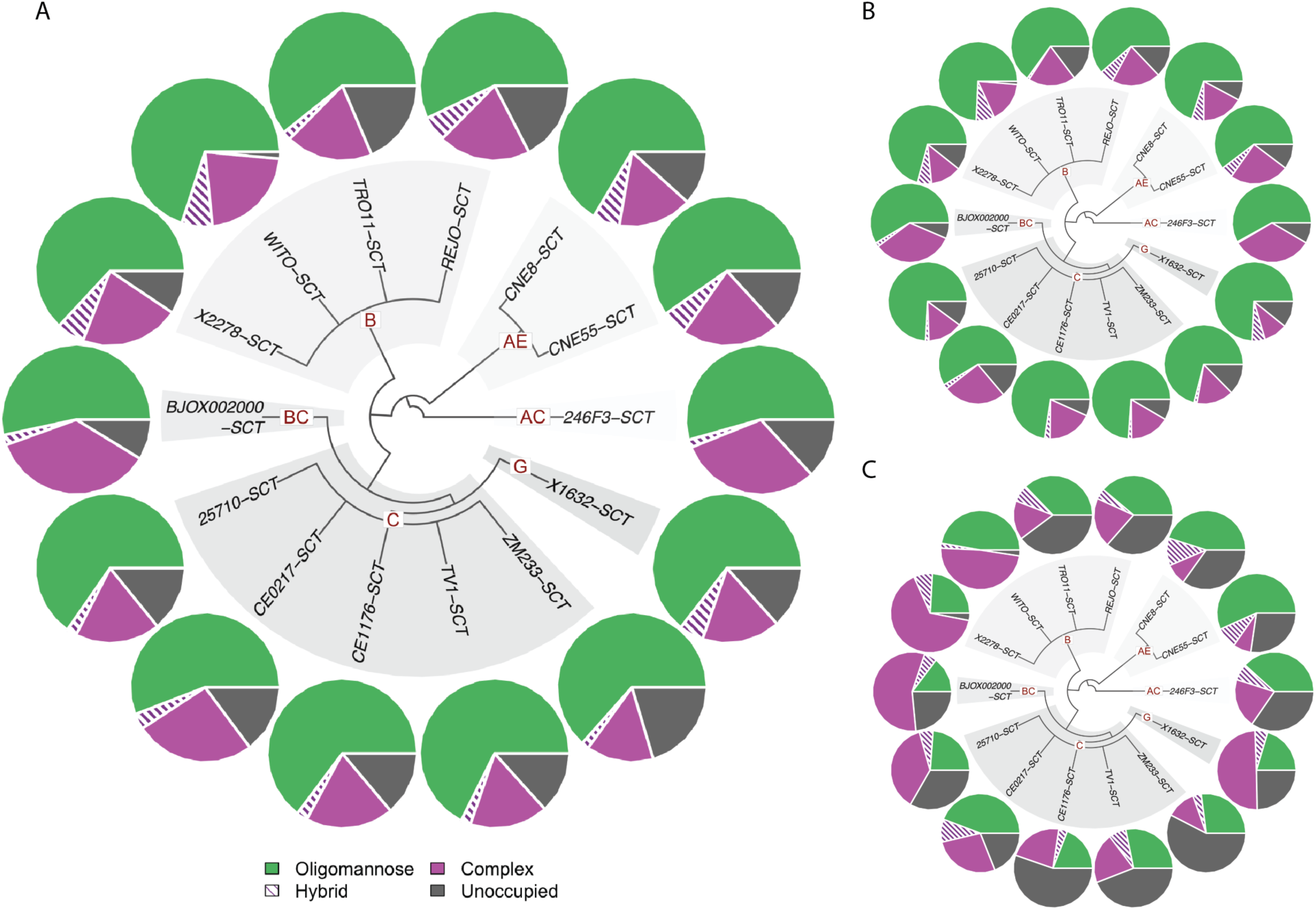
Site-specific glycan analysis of global panel SCT-stabilized HIV-1 trimers. **A.** Pie plots summarizing the overall glycan composition and site occupancy for the global panel of SCT-stabilized HIV-1 Env trimers. Only glycosylation sites that were reliably resolved by mass spectrometry are included. The plots reflect the relative proportions of oligomannose, complex, hybrid and unoccupied glycans across the full trimer. **B.** Pie plots representing the glycan composition and occupancy within the gp120 subunit of the global SCT panel. The gp120 region, which is more exposed and glycan-dense, shows a mix of high-mannose and complex-type glycans, reflective of typical HIV-1 Env glycosylation patterns and consistent with native-like folding. **C.** Pie plots representing the glycan composition and occupancy within the gp41 subunit of the SCT-stabilized trimers. The analysis highlights consistent glycosylation patterns across strains, with domain-specific differences in glycan processing and site occupancy.

In contrast to gp41, gp120 subunits were mostly occupied by oligonamannose, ranging from 58.5% (BJOX002000 SCT) to 74.3% (WITO SCT and X1632 SCT) (Figure 7B). LC-MS analysis of the relative abundance of oligomannose-type glycans at each site across all the samples analysed, revealed conservation of the intrinsic mannose patch (IMP), which is a feature of Env gp120 regardless of its quaternary context. IMP sites such as N262, N295, N332, N339, N363, N392 and N448 displayed predominantly oligomannose-type glycans in nearly all the single chain trimers (Figure S8). Additionally, N-linked glycan sites that comprise the trimer-associated mannose patch, which includes sites that are located along the protomer interface, such as N156 and N160, (key residues forming the epitope for V2-apex targeting bnAbs), were predominantly oligomannose-type glycans which is potentially indicative of efficient trimer formation (Figure S8). We did not observe significant differences between various clades, suggesting that the SCT design is a promising universal design to stabilize prefusion-like HIV trimers in the context of glycan occupancy.

### The SCT design is broadly applicable to diverse HIV Env strains, including Env lineage-based trimers known to elicit bnAbs in humans

To further validate the versatility of the SCT design strategy, we expanded our efforts beyond the global Env panel to include a larger set of HIV-1 Envs commonly used in vaccine development. This extended panel included Env lineage-based immunogens known to initiate or mature broadly neutralizing antibody (bnAb) responses in humans (Figure 8A, Table S3). All SCT constructs were expressed and purified via GNL affinity chromatography and analyzed by size-exclusion chromatography, which produced yields varying from 2.8 mg to 25 mg per liter 293F transfection (Table S3). Each construct exhibited a predominant trimeric peak, with moderate levels of aggregation and minimal unassembled monomers, indicating successful trimer assembly across the panel (Figure 8A). Antigenicity profiling using BLI confirmed that these SCTs maintain a native-like trimer structure across diverse subtypes. The majority of constructs demonstrated strong binding to bnAbs and low binding to non-neutralizing antibodies, consistent with a properly folded prefusion conformation (Figure 8B). This represents a major step forward, as previous stabilization strategies have shown limited applicability across diverse HIV-1 strains with vaccine potential.

**Figure 8.**
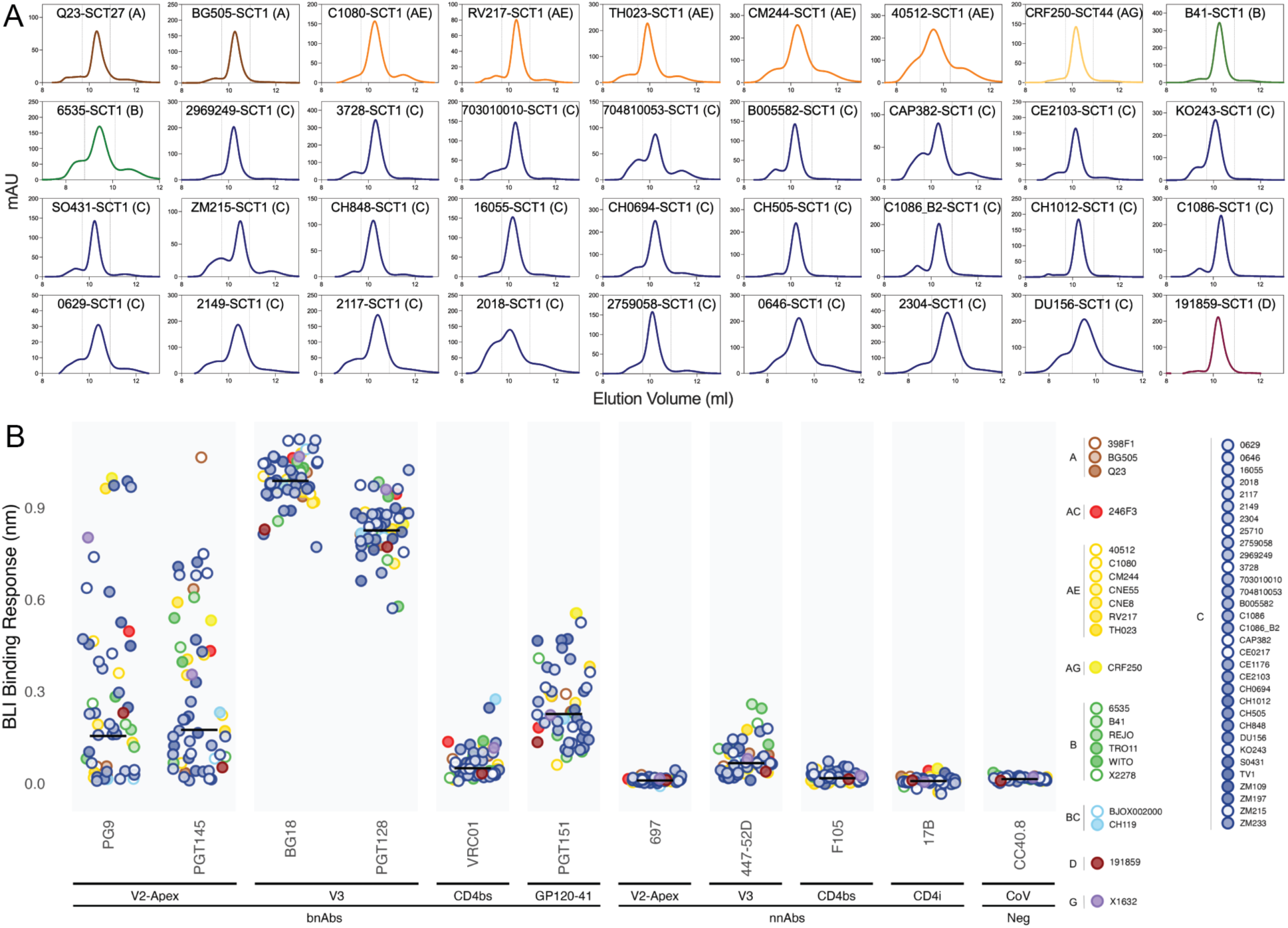
Design and characterization of a large panel of SCT-stabilized HIV-1 Env trimers. **A.** Size-exclusion chromatography (SEC) profiles of GNL-purified SCTs from an expanded panel representing diverse HIV-1 subtypes, analyzed on Superdex 200 columns. Each curve is color-coded to indicate a distinct HIV-1 subtype. Trimeric species elute within the region marked by dashed lines. A single, sharp peak in this region indicates consistent and proper trimer folding across the panel. B. Scatter plot showing BLI maximum binding responses of a representative panel of HIV bnAbs and non-neutralizing antibodies (nnAbs) to the GNL-purified SCTs (100 nM). The antibodies target a range of Env epitopes, allowing for assessment of antigenic integrity and epitope presentation across the large SCT panel.

In summary, the successful stabilization and antigenic validation of this expanded panel of SCTs, including lineage-based Envs, demonstrate the broad utility of the SCT platform. These constructs can serve as prime or boost immunogens in various vaccine regimens, including those designed to guide the induction of bnAbs through lineage-based strategies. Their compatibility with multiple delivery platforms, including mRNA, further underscores their potential to support next-generation HIV vaccine design aimed at eliciting protective broadly neutralizing antibody responses.

## Discussion

Eliciting broadly neutralizing antibodies remains a central goal of HIV vaccine development, yet the inherent instability and structural heterogeneity of the HIV Env trimer pose major challenges for immunogen design. Here, we present a generalizable and robust Single-Chain Trimer design platform that enables the stabilization and high-level expression of diverse HIV-1 Env trimers in their native-like, prefusion conformation. Our design integrates and builds upon previous stabilization strategies to yield soluble trimers with favorable antigenic profiles, structural integrity, and most importantly, compatibility with multiple vaccine delivery platforms. This SCT strategy offers a major advancement toward the development of HIV vaccines capable of guiding the induction of protective bnAb responses.

A key finding of this study is the broad applicability of the SCT platform across diverse HIV-1 strains. Beginning with Env-Ab lineage-based immunogens derived from the CAP256 donor, our iterative structure-guided design approach yielded the CAP256.SU-SCT9 construct, which demonstrated superior trimer integrity as revealed by a high-resolution cryo-EM structure, native-like antigenicity, and favorable glycan processing. Importantly, this design strategy was successfully extended to additional CAP256 Env lineage member as well as panels of HIV Envs representing global subtype diversity. SEC analysis, BLI profiling, and negative-stain EM consistently confirmed that these SCTs predominantly adopt prefusion-closed conformations with minimal aggregation or misfolding. Antigenicity studies revealed strong binding to trimer-dependent bnAbs and minimal recognition by non-neutralizing antibodies, reinforcing the ability of SCT design to present authentic neutralizing epitopes while masking irrelevant antigenic epitope surfaces.

Beyond structural and antigenic fidelity, our results emphasize the SCT platform’s compatibility with advanced vaccine delivery modalities. CAP256.SU-SCT9 exhibited highly favorable antigenicity when expressed on the cell surface, highlighting its potential as an immunogen for nucleic acid-based vaccine approaches such as mRNA, self-amplifying RNA (saRNA), and DNA. Notably, the binding profiles of membrane-bound SCT9 correlated strongly with those of the soluble trimer and with pseudovirus neutralization titers, validating its functional and structural authenticity in both soluble and membrane-anchored formats. This finding is particularly relevant given the growing prominence of mRNA-based vaccines, where proper folding and antigen display in vivo are essential (Chaudhary et al., 2021; Gote et al., 2023). Additionally, the SCT design proved readily adaptable for multivalent display via self-assembling nanoparticle platforms. Grafting SCT9 onto a ferritin scaffold generated uniformly assembled nanoparticles displaying eight trimer copies per particle. Homogeneous assembly has been shown to be critical for vaccine efficacy, whereas multimerized HIV-1 immunogens with heterogeneous assembly result in suboptimal immune responses (Mu et al., 2024). These nanoparticles exhibited enhanced binding to multiple bnAbs, including precursor V2-apex bnAbs such as CAP256.UCA, consistent with avidity-driven epitope presentation. Negative-stain EM confirmed the structural homogeneity of the trimer nanoparticles, supporting their potential use as priming immunogens for initiating rare bnAb lineages.

Glycan profiling of the SCT panel further validated the native-like properties of the constructs. LC-MS analysis revealed a predominance of oligomannose-type glycans on gp120 and conservation of key glycan patches such as the intrinsic mannose patch (IMP) and the trimer-associated mannose patch (TAMP). These glycan signatures are characteristic of properly folded native Env trimers and are essential for accurate epitope presentation and bnAb recognition (Behrens *et al*., 2016; Cao et al., 2017; Seabright et al., 2019). The consistent glycosylation patterns observed across multiple HIV-1 subtypes and clades support the SCT design as a reliable platform for reproducing native-like glycan profiles.

Finally, the successful stabilization of multiple lineage-based Envs known to elicit bnAbs in humans underscores the relevance of SCT design platform for next-generation vaccine strategies. These constructs can be incorporated into Env lineage-based, or germline-targeting prime-boost regimens designed to shepherd bnAb precursors through affinity maturation. The SCT design’s structural integrity, antigenic fidelity, and delivery flexibility positions it as a versatile platform to expand the immunogen repertoire required for these complex multi-stage approaches (Andrabi *et al*., 2018).

In summary, our findings establish the SCT design as a powerful and generalizable strategy for engineering native-like HIV Env trimers. By preserving key structural and antigenic features across a broad spectrum of HIV-1 strains, and enabling delivery through mRNA, DNA, and nanoparticle platforms, this approach supports the rational development of diverse immunogen regimens for bnAb induction. As HIV vaccine research continues to advance through iterations in immunogen design and reverse vaccinology, the SCT platform offers a foundation for achieving the long-sought goal of a protective, broadly effective HIV vaccine.

## Resource Availability

### Lead contact

Further information and requests for resources and reagents should be directed to and will be fulfilled by the Lead Contact, Raiees Andrabi

### Materials availability

Upon specific request and execution of a material transfer agreement (MTA) from The University of Pennsylvania to the Lead Contact, DNA plasmids will be made available.

### Data and code availability

The data supporting the findings of this study are available within the published article and summarized in the corresponding tables, figures, and supplemental materials. The single chain trimer sequences have been deposited in GenBank under accession numbers XXX-YYY. The atomic models and cryo-EM maps generated for the CAP256SU.SCT9 – 40591-a.05 complex have been deposited to the Protein Data Bank (PDB; http://www.rcsb.org/) and the Electron Microscopy Databank (EMDB; http://www.emdataresource.org/) under accession codes PDB 9YU0 and EMD 73498, respectively. The atomic model and corresponding structure factor file for the 40591-a.05 Fab have been deposited in the PDB under accession code 9YU1.

## Acknowledgments

This work was supported by National Institute of Allergy and Infectious Diseases National Institutes of Health grants R01 AI 167716 (R.A.), R61 AI 161818 (R.A., G.M.S.), U19 AI 188562 (R.A.), P01 AI 177683 (R.A.) and UM1 AI144462 (A.B.W.). The content is solely the responsibility of the authors and does not necessarily represent the official views of the National Institutes of Health. This study was made possible by the generous support of the Bill and Melinda Gates Foundation through the Collaboration for AIDS Vaccine Discovery (CAVD) grants INV041767 and INV064777 (G.M.S., and R.A.) This study was supported in part by funds from the Howard Hughes Medical Institute Emerging Pathogens Initiative (to C.O.B.). Additionally, C.O.B. is supported by the Howard Hughes Medical Institute Hanna Gray Fellowship, Rita Allen Foundation, Pew Biomedical Scholars Program, and is a Chan Zuckerberg Biohub investigator. The work in this study was additionally supported by funds from the Gates Foundation through the collaboration for AIDS Vaccine Discovery grant INV/070116 (M.C). Cryo-EM data for this work was collected at the Stanford-SLAC cryo-EM resource center with support from Haoqing Wang and Bharti Singal. X-ray crystallographic data were collected at the Stanford Synchrotron Radiation Lightsource, SLAC National Accelerator Laboratory, which is supported by the U.S. Department of Energy, Office of Science, Office of Basic Energy Sciences under Contract No. DE-AC02-76SF00515. The SSRL Structural Molecular Biology Program is supported by the DOE Office of Biological and Environmental Research, and by the National Institutes of Health, National Institute of General Medical Sciences (P30GM133894). We thank Daniel Fernandez and Beamline 12-2 staff for their support.

## Author Contributions

X.L., N.M., R.A. conceived and designed experiments; S.C., W.H. designed self-assembling nanoparticle platform; X.L., N.M., S.C., A.S., G.A., K.A., T.S., P.O., Y.Z., B.L., G.G., A.L.V., S.A.,I.J., A.M., T.C., F.A. conducted HIV trimer and antibodies transfection and purification, size exclusion chromatography; X.L., R.R.C., T.S. performed cell surface binding assay; X.L., N.M., A.S., G.A. performed soluble trimer BLI binding assay; X.L., A.S., R.N. performed neutralization assay; I.K., J.L.T., W.L., G.O., A.B.W., C.O.B. performed nsEM; I.K., C.O.B performed cryo-EM high-resolution structures of antibodies in complex with trimers and structural analysis with contributions from R.H., F.B.R., B.H.H., D.R.B., G.M.S.; N.E.J., J.D.A., M.C performed glycan analysis; X.L., I.K., N.E.J., S.D.S., C.O.B., R.A. wrote the manuscript with input from all listed authors. All authors were involved in critical revision of the manuscript for accuracy and important intellectual content.

## Material and Methods

### HIV Envelope single chain trimer protein design

For the design of the SCT constructs we purchased g-blocks (IDT) encoding for the Env region, with each containing specific SCT mutations. We preformed Gibson Assembly to clone the Env fragments into expression vectors (Genlantis) following the manufacturer’s protocol, and successful incorporation of mutations was verified through sequencing analysis (Azenta).

### Cell Lines

Expi293F cells (Gibco, Cat# A14527) were cultured in Expi293 Expression Medium (Gibco, Cat# A1435101) at 37°C with 8% CO₂ in a 125rpm shaker. HEK293F cells (Gibco, Cat# A14527) were cultured in Freestyle medium (Gibco, Cat# 12338-018) at 37°C with 8% CO₂ in a 125rpm shaker. HEK293T cells (ATCC, Cat# CRL-3216) were maintained in Dulbecco’s Modified Eagle Medium (DMEM) (Corning, Cat# 10-017-CV) supplemented with 10% heat-inactivated fetal bovine serum (FBS) (Thermo Fisher, Cat# MT35016CV), 4 mM L-glutamine (Corning, Cat# 25-005-CI), and 1% penicillin-streptomycin (P/S) (Corning, Cat# 30-002-CI) at 37°C in a 5% CO₂ incubator. TZM-bl 931 cells (NIH AIDS Reagents Program) were used for the pseudovirus neutralization assay.

### Expression and purification of HIV single chain trimer

Plasmids encoding the Env trimers were transfected into HEK293F cells using OptiMEM and PEI-MAX 40K transfection reagent (Kyfora, cat# 24765-1). The cells were incubated, and four days post-transfection, the supernatants containing the expressed protein trimers were collected. Purification was carried out using affinity chromatography with either agarose-bound Galanthus nivalis lectin (GNL) (Vector Labs, cat #AL-1243-5) or TOYOPEARL AF-Tresyl-650M beads (TOSOH, cat #0014472) conjugated to the PGT145 broadly neutralizing antibody (bnAb). To ensure further purification, the eluates were subjected to size-exclusion chromatography (SEC) using a Superdex 200 Increase 10/300 GL column (GE Healthcare, cat #GE28-9909-44) and Superose 6 Increase 10/300 GL columns (Cytiva) in PBS.

### Expression and purification of HIV bnAb and nnAb

Paired heavy and light chain plasmids for given HIV bnAb and nnAb were co-transfected in Expi293 cells (Thermo Fisher Scientific, cat #A14527) in a 1:1 ratio using FectoPRO (Polyplus, cat #116-001) transfection reagent and were fed with 0.3M valproic acid (Sigma, cat #P4543-100G) and 40% Glucose (Gibco, #A2494001) 24 hours after the transfection. Monoclonal IgGs were purified from the culture supernatant five days post-transfection with 1:1 ratio Protein-A (Cytiva, cat #17127903) and Protein-G Sepharose beads (Cytiva, cat #17061805) per manufacturer’s instructions. After elution with IgG elution buffer (Thermo Fisher Scientific, cat #PI21009), antibodies were buffer exchanged into PBS using 50kDa Ultra centrifugal filter unit. (Millipore, cat #UFC905024)

### Neutralization assay

Monoclonal HIV antibodies (bnAbs, nnAbs) (mAbs, 10μg/ml) were 5-fold serially diluted in complete DMEM (cDMEM) (25ul) and incubated with HIV-1 Env-pseudotyped virus (25μl) for 60 minutes at 37°C in duplicate 96-well Culture Plates. TZM-bl cells (20,000 cells per well) with 40 μg/ml DEAE-Dextran were then added (50μl) and incubated overnight. Control wells included cells only (background) and virus only (maximal entry). Serial dilutions were performed with tip changes to prevent carryover. Luciferase activity was measured using the Bright-Glo Luciferase Reporter Assay (Promega) and a SpectraMax L luminometer. Percent neutralization was calculated as: ((RLU_Virus_ – RLU_test_) / RLU_Virus_) x 100. Background RLU from uninfected control wells was subtracted before final calculations. Neutralizing antibody titers (IC₅₀) were determined via a four-parameter nonlinear dose-response curve.

### **BioLayer** Interferometry **(BLI)**

For high throughput antigenicity screening of HIV Single Chain Trimer designs, BLI was performed with HIV bnAbs and nnAbs (10 µg/ml). For screening, IgGs were immobilized on ProA sensors (Sartorius) to a signal of at least 1.0 nm using an Octet Red96 instrument (ForteBio). The immobilized IgGs were then dipped in the running buffer (PBS, 0.1% BSA, 0.02% Tween20, pH 7.4). Following a 120 s association period with 100 or 500 nM of designed SCTs, the tips were dipped into the running buffer and dissociation was measured for 240 s.

### Cell surface binding assay (CELISA)

HEK293T cells (7 x10^6^) were seeded in T175 flasks one day prior to transfection with plasmids encoding the Single Chain Trimers with transmembrane domain using Lipofectamine 2000 (Thermo Fisher Scientific, cat #11668500), following the manufacturer’s protocol. After 48 hours of incubation at 37°C with 5% CO₂, cells were harvested using FACS buffer (PBS + 2% FBS+5mM EDTA (Invitrogen, cat #15575-038). 0.1x10^6^ cells were aliquoted into U bottom 96-well plates followed by centrifugation for 1500 rpm for 2 mins. Supernatant was removed and antibodies diluted in FACS buffer (10ug/ml) were added to wells with 2-fold serial dilution. After 1 hour incubation at 4°C, cells were washed two times with FACS buffer and subsequently stained with Mouse Anti-Human IgG FC-PE (SouthernBiotech, cat #9040-09) for 1 hour at 4°C in the dark. Finally, cells were washed three times and resuspended in FACS buffer before running on a Bio-Rad ZE5 flow cytometer. Data were analyzed using FlowJo software.

### Site-specific glycan analysis of HIV SCTs

50 µg aliquots of each sample were denatured for 1h in 50 mM Tris/HCl, pH 8.0 containing 6 M of urea and 5 mM dithiothreitol (DTT). Next, Env samples were reduced and alkylated by adding 20 mM iodoacetamide (IAA) and incubated for 1h in the dark, followed by a 1h incubation with 20 mM DTT to eliminate residual IAA. The alkylated Env samples were buffer exchanged into 50 mM Tris/HCl, pH 8.0 using Vivaspin columns (10 kDa) and aliquots were digested separately overnight using Trypsin (Mass Spectrometry Grade, Promega), chymotrypsin (Promega), or alpha-lytic protease (Sigma Aldrich) at a ratio of 1:30 (w/w). The next day, the peptides were dried and extracted using an Oasis HLB µElution Plate (Waters).

The peptides were dried again, re-suspended in 0.1% formic acid, and analyzed by nanoLC-ESI MS with a Vanquish Neo (Thermo Fisher Scientific) system coupled to an Orbitrap Eclipse Tribrid mass spectrometer (Thermo Fisher Scientific) using stepped higher energy collision-induced dissociation (HCD) fragmentation. Peptides were separated using a μPAC™ Neo HPLC Column (180 µm × 110 cm). A trapping column (PepMap 100 C18 3μM 75μM × 2cm) was used in line with the LC prior to separation with the analytical column. The LC conditions were as follows: 280-minute linear gradient consisting of 4-32% acetonitrile in 0.1% formic acid over 260 minutes followed by 20 minutes of alternating 76% acetonitrile in 0.1% formic acid and 4% ACN in 0.1% formic acid, used to ensure all the sample had eluted from the column. The flow rate was set to 300 nL/min. The spray voltage was set to 2.5 kV and the temperature of the heated capillary was set to 55 °C. The ion transfer tube temperature was set to 275 °C. The scan range was 350−2000 m/z. The stepped HCD collision energies were set to 15, 25 and 45% and the MS2 for each energy was combined. Precursor and fragment detection was performed using an Orbitrap at a resolution MS^1^ = 120,000, MS^2^ = 30,000. A standard AGC target for MS^1^ (4e^5^) and MS^2^ (1e^4^) and auto injection times (MS^1^ =50ms MS^2^ =54ms) were used.

Glycopeptide fragmentation data were extracted from the raw file using Byos (Version 5.5; Protein Metrics Inc.). The glycopeptide fragmentation data were evaluated manually for each glycopeptide; the peptide was scored as true-positive when the correct b and y fragment ions were observed along with oxonium ions corresponding to the glycan identified. The MS data was searched using the Protein Metrics 305 N-glycan library. The relative amounts of each glycan at each site as well as the unoccupied proportion were determined by comparing the extracted chromatographic areas for different glycotypes with an identical peptide sequence. All charge states for a single glycopeptide were summed. The precursor mass tolerance was set at 4 ppm and 10 ppm for fragments. A 1% false discovery rate (FDR) was applied. The relative amounts of each glycan at each site which yielded data as well as the unoccupied proportion were determined by comparing the extracted ion chromatographic areas for different glycopeptides with an identical peptide sequence. Glycans were categorized according to the composition detected.

HexNAc(2)Hex(9−3) was classified as M9 to M3. Any of these structures containing a fucose were categorised as FM (fucosylated mannose). Complex-type glycans were classified according to the number of HexNAc subunits and the presence or absence of fucose. Core glycans refer to truncated structures smaller than M3. As this fragmentation method does not provide linkage information, compositional isomers are grouped.

To obtain data for sites that frequently present low intensity glycopeptides, the glycans present on the glycopeptides were homogenized to boost the intensity of these peptides. The remaining glycopeptides were first digested with Endo H (New England Biolabs) to deplete oligomannose- and hybrid-type glycans and leave a single GlcNAc or GlcNAcFuc residue at the corresponding site. The reaction mixture was then dried completely and resuspended in a mixture containing 50 mM ammonium bicarbonate and PNGase F (New England Biolabs) using only H_2_O^18^ (Sigma-Aldrich) throughout. The resultant peptides were purified and subjected to reverse-phase (RP) nanoLC-MS. Three modifications were searched for: +203 Da corresponding to a single GlcNAc, or +349 corresponding to a GlcNAcFuc, a remnant of an oligomannose/hybrid glycan (+/-fucose), and +3 Da corresponding to the O^18^ deamidation product of a complex glycan. Data acquisition and analysis was performed as above and the relative amounts of each glycoform were determined, including unoccupied peptides. Not all PNGS present in each protein could be resolved.

### X-ray crystallography

40591-a.05 Fab in Tris-buffered saline (TBS, 20 mM Tris pH 8.0, 150 mM NaCl) was concentrated to 15 mg/mL using an Amicon spin filter with a 30-kDa molecular weight cutoff (Millipore-Sigma). Crystals were set up using the sitting drop vapor diffusion method and equal volumes of 40591-a.05 Fab and reservoir were mixed using a Mosquito LCP liquid handling robot (SPT LabTech) and commercially available 96-well crystallization screens (Hampton Research). Crystals were grown at room temperature and observed under multiple conditions. The single crystal used for structure determination was obtained in 0.2 M ammonium tartrate and 30% PEG600 (v/v), and cryoprotected in a solution matching the reservoir and 30% glycerol prior to cryocooling in liquid nitrogen.

X-ray diffraction data was collected at the Stanford Synchrotron Radiation Lightsource (SSRL) beamline 12-2 with an Eiger X 16M pixel detector (Dectris) at a wavelength of 0.979 Å and temperature of 100 K. Diffraction data from a single crystal were indexed and integrated in X-ray Detector Software (XDS) (Kabsch, 2010) and then merged using AIMLESS in CCP4 (Agirre et al., 2023). Structures were determined using molecular replacement in PHASER,(McCoy et al., 2007) using the following starting models for the V_H_ (PDB: 6CNR) and V_L_ (PDB: 6PHG), with the CDRH_3_ and CDRL_3_ trimmed, respectively. Coordinates were subsequently refined using iterative rounds of automated and manual refinement in Phenix (Zwart et al., 2008) and Coot (Emsley et al., 2010), respectively. The final model contains 98.61% Ramachandran-favored outliers, 0.84% rotamer outliers, and 0% Ramachandran outliers. All statistics for the final model can be found in the Supplementary Table S2.

### Cryo-EM sample preparation

Quantifoil Cu 1.2/1.3 300 mesh grids (Electron Microscopy Sciences) were glow discharged for 90 s at 10 mA (PELCO easiGlow) before sample application. 2 mg of purified SCT9 (12.2 mg/mL) were incubated with a 3x molar excess of 40591-a.05 Fab overnight at room temperature in TBS with 0.01% sodium azide. Complexes were purified via size-exclusion chromatography (SEC) using a Superose 6 10/300 Increase column (GE Healthcare) in 1X TBS with 0.01% sodium azide to remove the excess Fab. Fractions corresponding to the CAP256SU.SCT9 – 40591-a.05 complex were pooled and concentrated. Fluorinated octyl-maltoside (Anatrace) was added to the sample to a final concentration of 0.025% w/v and 3 uL of complex was applied to each grid twice, with manual blotting in between to achieve adequate particle concentration. A Mark IV Vitrobot (Thermofisher) was used to blot grids for 3 s with a blot force of 3 at 7°C and 100% humidity. Grids were plunge frozen and vitrified in liquid ethane and stored in liquid nitrogen until screening and data acquisition.

### Cryo-EM data collection

Single-particle cryo-EM datasets were collected on a 300 kV Titan Krios transmission electron microscope (Thermo Fisher Scientific) equipped with a Falcon 4i direct electron detector and an energy filter (Selectris, 10 eV slit width). Movies were acquired in an automated fashion using EPU (Thermofisher) and a multishot data acquisition pattern, with a nominal magnification of 130,000 x, and a calibrated pixel size of 0.92 Å, with a total dose of 40 e/Å^2^ and a defocus range of -1 to -2.4 µm.

### Cryo-EM data processing

For all data sets, motion correction, contrast transfer function (CTF) estimation, and reference-free particle picking were performed using cryoSPARC v4.6.2 (Punjani et al., 2017). Micrographs with ice contamination, CTF-estimated resolutions worse than 7 Å and ice thickness greater than a score of 1.25 were discarded. 2D classes were generated and a subset of down-sampled, reference-free particles from the 2D classes were used to generate ab-initio reconstructions in cryoSPARC. Selected 2D classes of reference-free particles were used as references for template-based particle picking and were used for all subsequent processing. Heterogeneous refinement of particles was performed to generate an initial reference model. Iterative 3D classification was performed in cryoSPARC, alternating between homogeneous and heterogeneous refinement, followed by re-extraction without binning and nonuniform refinement in cryoSPARC. Further details regarding data collection and data processing workflows can be found in the supplementary information.

### Cryo-EM model building and structure determination

Initial coordinates were determined using ModelAngelo to first fit the cryo-EM density from input sequence in an un-biased manner (Jamali et al., 2024). The model was further refined in PHENIX using real space refinement and manual building in Coot, based off the quality of the map (Emsley *et al*., 2010; Zwart *et al*., 2008). For certain residues, side chains were not included if sufficient density was not visible. Glycans were modeled at potential N-linked glycosylation sites using the carbohydrates module in Coot, when justified by the density. Model quality was evaluated using MolProbity (Chen et al., 2010). Trimer numbering was denoted by HXB2 numbering, and antibody residues were denoted by Kabat numbering.

### nsEM sample preparation

All samples (trimers or nanoparticles) were diluted to 20 ng/uL in 1X Tris-buffered saline. 3 uL of sample was applied to glow-discharged C-flat carbon grids and allowed to incubate on the grids for 1 minute 30 seconds before manual blotting. Grids were either washed three times with filtered deionized water and then washed twice with filtered 1% (w/v) uranyl formate or stained directly one time with 2% (w/v) uranyl formate without the wash steps. Staining was performed for 1 minute 30 seconds before manual blotting.

### nsEM data collection and processing

nSEM datasets were collected on one of three microscope configurations: 1) 200 kV Glacios transmission electron microscope (Thermo Fisher Scientific) equipped with a Falcon 4i direct electron detector. (Thermo Fisher Scientific), 2) 200 kV Talos transmission electron microscopy (Thermo Fisher Scientific) equipped with a CETA 4K CMOS (Thermo Fisher Scientific), or 3) 120 kV Tecnai Spirit (FEI) equipped with an Eagle 4K CCD (Thermo Fisher Scientific). Glacios m. icrographs were acquired in an automated fashion using EPU (Thermofisher) using a generated pattern with images taken 2.8 µm apart, with a nominal magnification of 57,000 x, and a calibrated pixel size of 2.5 Å, with a total dose of 15 e/Å^2^ and an applied defocus of -2 µm. Talos micrographs were collected using Leginon (Cheng et al., 2021) with a nominal magnification of 73,000 x and a calibrated pixel size of 1.98 Å with a total dose of 25 e/Å^2^ and applied defocus of -1.5 µm. Tecnai Spirit micrographs were collected using Leginon with a nominal magnification of 52,000 x and a calibrated pixel size of 2.06 Å with a total dose of 25 e/Å^2^ and applied defocus of -1.5 µm. For all data sets, contrast transfer function (CTF) estimation, and reference-free particle picking were performed using cryoSPARC v4.6. (Punjani *et al*., 2017). Reference-free particles were initially picked and 2D classification was performed. Selected 2D classes were subsequently used as references for template-based picking and further 2D classification.

### Statistical Analysis

Statistical analyses were performed in GraphPad Prism and R. Data were considered statistically significant when p < 0.05.

## Supplementary Figures and Tables

**Figure S1.**
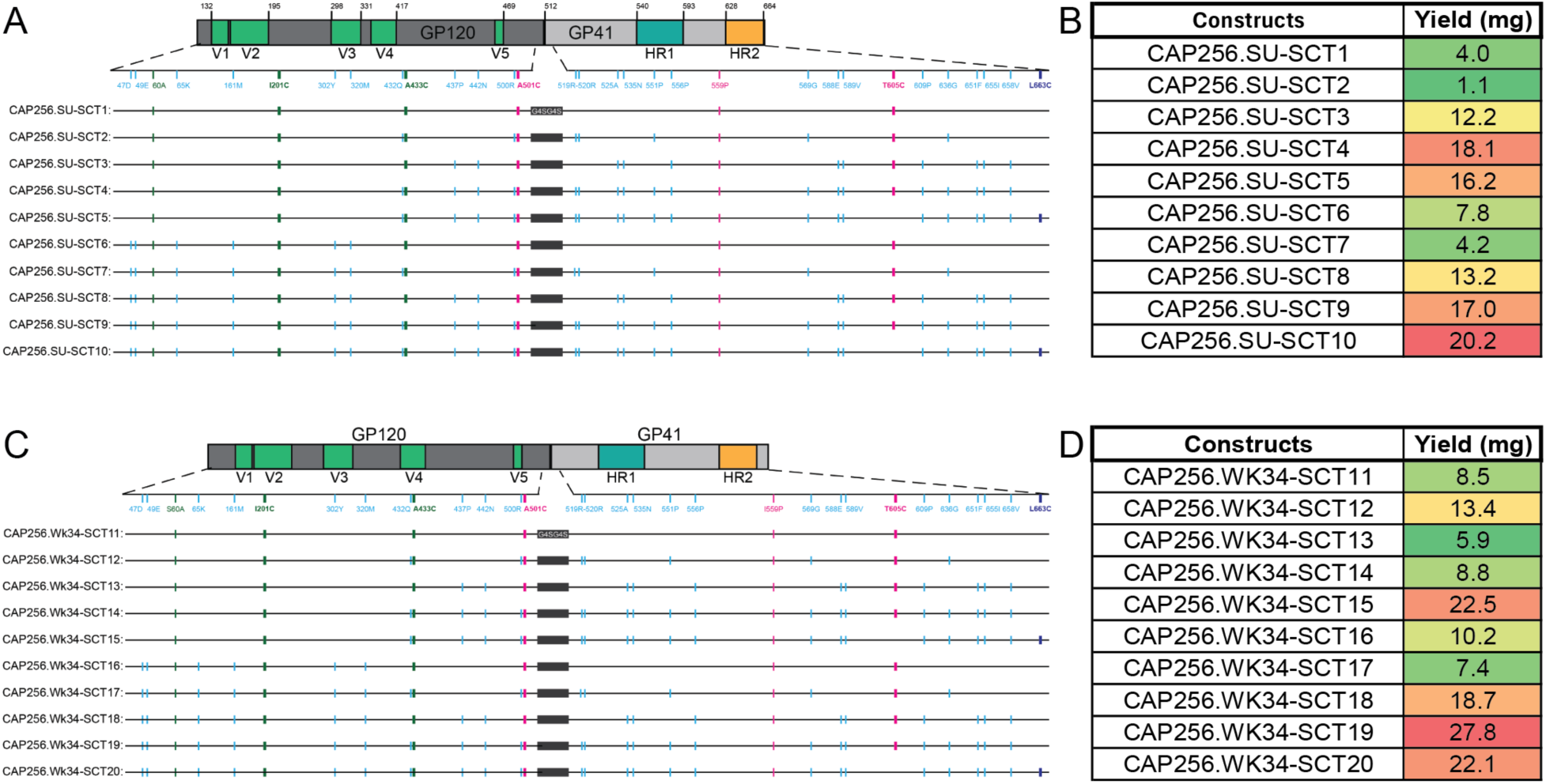
Design and Expression Yields of CAP256.SU and CAP256.Wk34 SCT-Stabilized HIV-1 Env Trimers (SCT1–20) **A.** Schematic sequence alignment of the gp120 and gp41 subunits from the CAP256.SU-SCT1– 10 constructs, highlighting all introduced mutations. Mutations are color-coded based on the category of stabilization: NFL (native flexibly linked) in green, RnS (repair-and-stabilized) in blue, and DS-SOSIP (disulfide-stabilized SOSIP) in pink. This alignment illustrates the progressive engineering applied to optimize trimer stability and antigenicity. **B.** Table summarizing the protein yields (mg/L) of the CAP256.SU-SCT1–10 constructs following transient transfection of 293F cells and purification using Galanthus nivalis lectin (GNL) affinity chromatography. Yield variations reflect construct-specific effects on expression and folding efficiency. **C.** Schematic sequence alignment of the gp120 and gp41 subunits for the CAP256.Wk34-SCT11–20 constructs, with all engineered mutations indicated. Color coding follows the same scheme as in panel A (NFL in green, RnS in blue, DS-SOSIP in pink), illustrating the application of diverse stabilizing strategies across this second set of Env trimers derived from the CAP256.Wk34 strain. **D.** Table summarizing the protein yields (mg/L) of the CAP256.Wk34-SCT11–20 constructs following GNL purification from 293F transfections (100 ml). These data provide a comparative overview of expression efficiency across the extended SCT panel.

**Figure S2.**
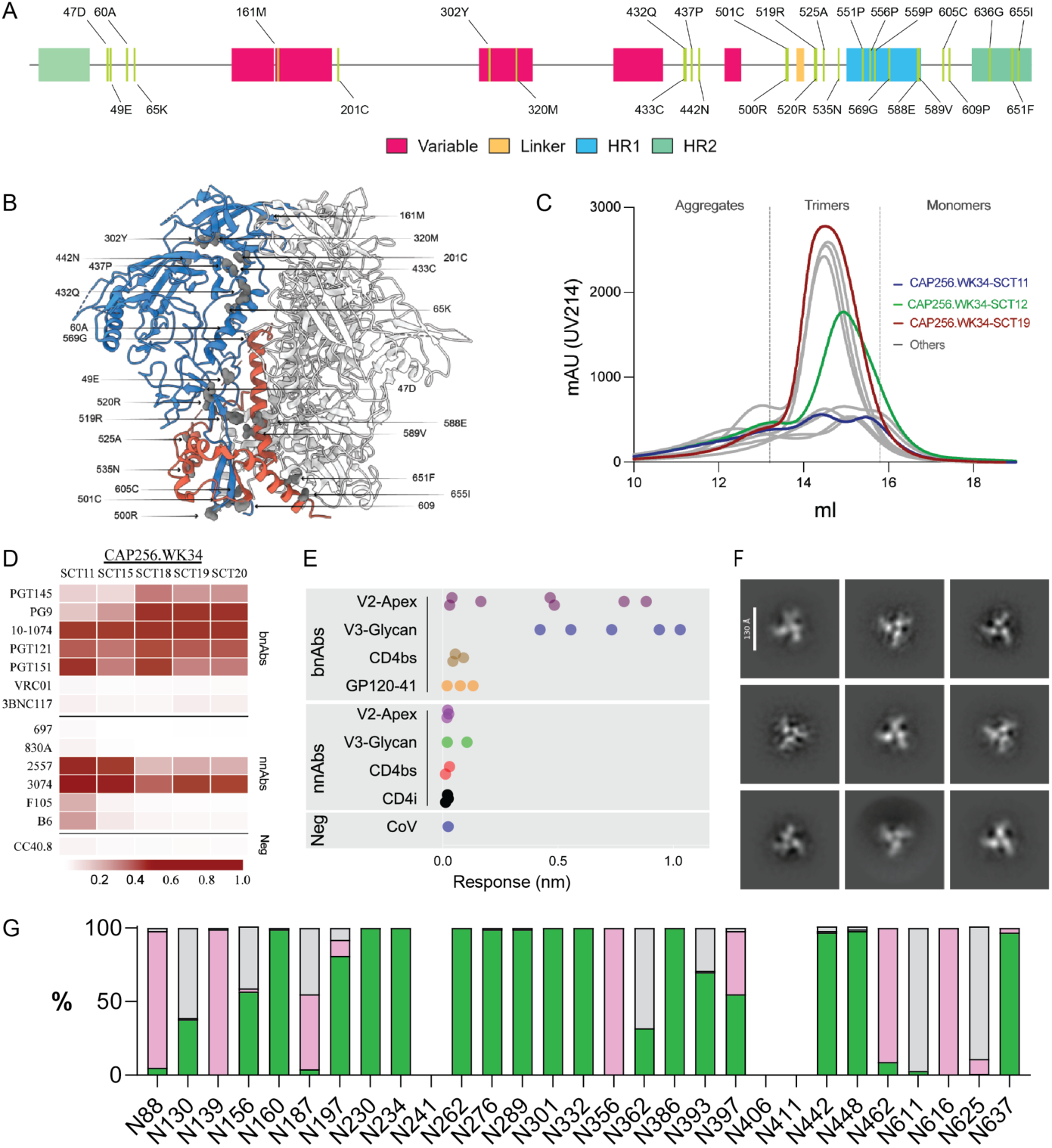
Design and characterization of a single-chain trimer (SCT) based on CAP256.Wk34 Env to enhance expression of native-like soluble trimers. **A.** Gene map illustrating point mutations incorporated into the SCT design. SCT-specific mutations are shown in grey. Heptad repeat regions HR1 and HR2 are highlighted in blue and green, respectively, alongside other distinct regions within the CAP256.Wk34 Env. **B.** Schematic representation of SCT mutations mapped onto the CAP256.Wk34 Env. The gp120 region is highlighted in blue, gp41 in red, and SCT mutations are indicated in grey. **C.** Size exclusion chromatography (SEC) profiles of CAP256.Wk34-SCT11 to SCT20 on a Superdex 200 column. The base construct CAP256.Wk34-DS-SOSIP (SCT11) is shown in blue, CAP256.Wk34-NFL (SCT12) in green, and CAP256.Wk34-SCT19 in red. Dashed lines indicate separation of trimeric forms from aggregated or dimeric/monomeric Env species. **D.** Heatmap showing the maximum BioLayer Interferometry (BLI) binding response of selected well-assembled CAP256.Wk34-SCTs (100 nM) to a panel of HIV-1 bnAbs and non-neutralizing antibodies (nnAbs). CC40.8, a SARS-CoV-2 stem-helix antibody, is included as a negative control. **E.** Dot plot displaying the maximum BLI binding response of CAP256.Wk34-SCT19 (100 nM) to an expanded panel of HIV-1 bnAbs and nnAbs. **F.** Negative-stain electron microscopy (nsEM) 2D class averages of CAP256.Wk34-SCT19 reveal that the trimers adopt a 100% native-like conformation. **G.** Glycan occupancy profile assessed by site-specific glycan analysis (SSGA) using proteomics. High-mannose glycans are shown in green, complex glycans in pink, and unoccupied sites in grey.

**Figure S3.**
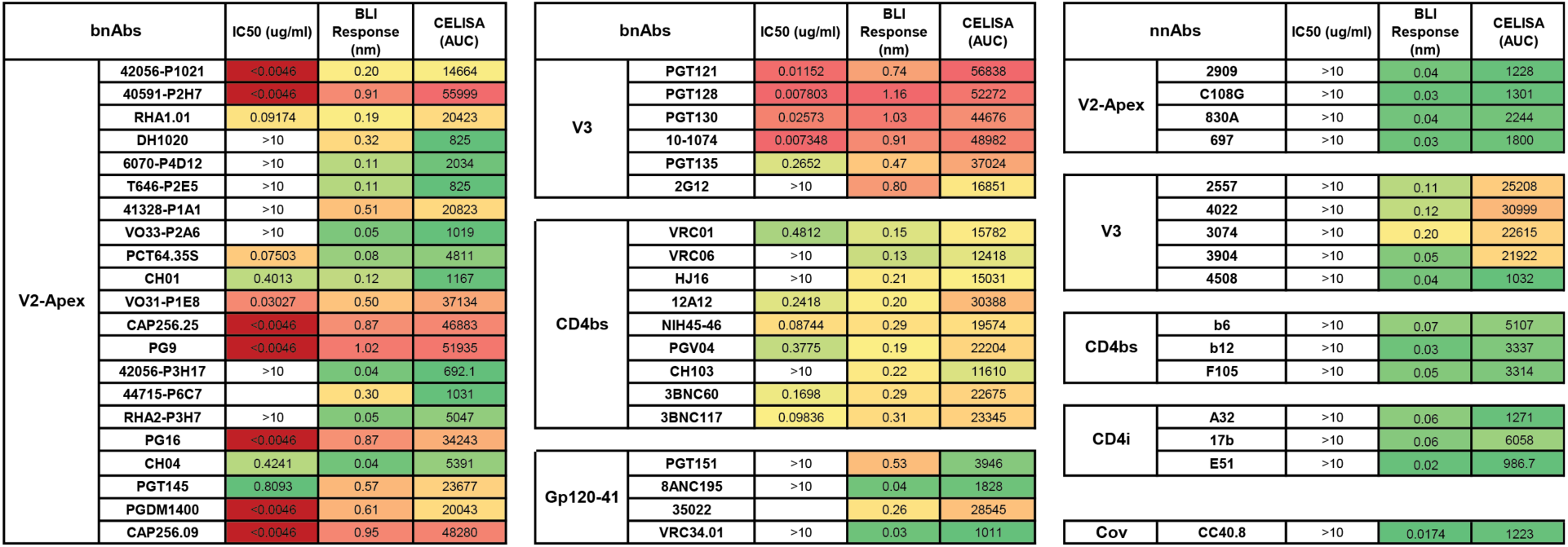
Antigenic profile of soluble and membrane-bound CAP256.SU-SCT9 trimers. Heatmap representing CAP256.SU Env-pseudotyped virus neutralization IC_50_ titers, biolayer Interferometry (BLI) maximum binding responses of the soluble CAP256.SU-SCT9 trimer (at 500 nM), and cell surface-expressed CAP256.SU-SCT9 trimer binding (CELISA) to a panel of HIV bnAbs and non-neutralizing antibodies (nnAbs) targeting diverse Env epitopes.

**Figure S4.**
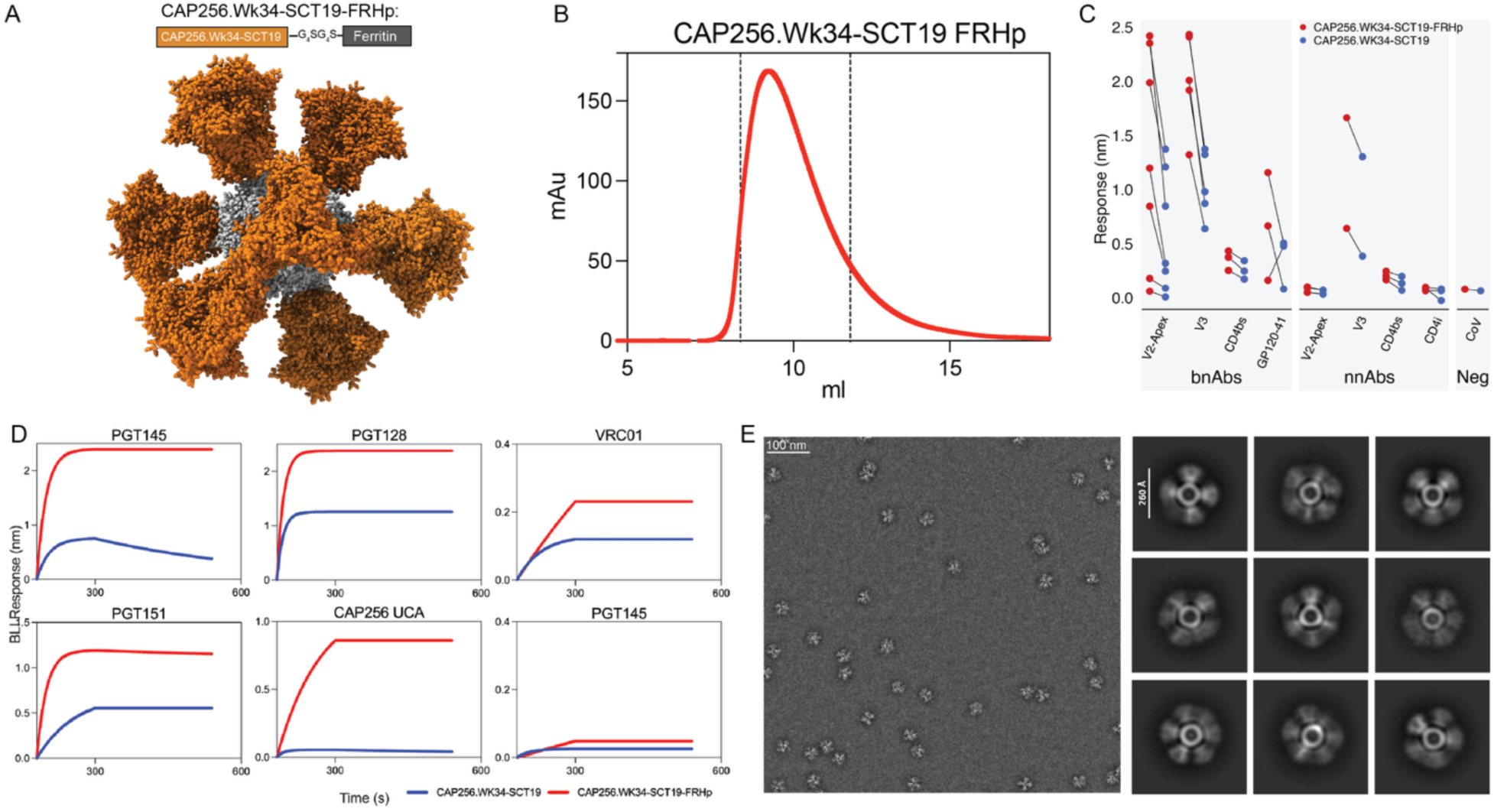
Design and characterization of self-assembled ferritin nanoparticles displaying CAP256.Wk34-SCT19 trimers for enhanced antigenicity. **A.** Schematic representation of the CAP256.Wk34-SCT19-FRHp ferritin nanoparticle design. Each nanoparticle displays 8 copies of the CAP256.Wk34-SCT19 trimer (highlighted in green) arranged on the surface of a 24-subunit ferritin core (shown in grey), allowing multivalent presentation of the Env trimer. **B.** Size exclusion chromatography (SEC) profile of the CAP256.Wk34-SCT19-FRHp nanoparticle, following purification by PGT145 affinity chromatography and subsequent SEC on a Superose6 column. A single nanoparticle peak eluting at ∼10 mL is shown within the dashed lines, indicating a uniform and stable nanoparticle population. **C.** Dot plot comparing the maximum BLI binding responses of the CAP256.Wk34-SCT19 soluble trimer (500 nM) and the CAP256.Wk34-SCT19-FRHp nanoparticle (500 nM) to a panel of HIV bnAbs and non-neutralizing antibodies (nnAbs) targeting diverse Env epitopes. Lines connect binding values for each antibody to both the trimer and the nanoparticle, highlighting differences in binding avidity. **D.** BLI binding kinetics (association and dissociation curves) comparing CAP256.Wk34-SCT19 trimer (500 nM, blue) and CAP256.Wk34-SCT19-FRHp nanoparticle (500 nM, red) to a representative set of HIV bnAbs, including V2-apex-targeting unmutated common ancestor (UCA) antibody, CAP256 UCA. CC40.8, a SARS-CoV-2 S2 stem-targeting antibody, serves as a negative control. The multivalent nanoparticle shows enhanced avidity binding and slower off-rates compared to the soluble trimer. **E.** Negative-stain electron microscopy (nsEM) of CAP256.Wk34-SCT19-FRHp nanoparticle. A representative micrograph (left) and corresponding 2D class averages (right) reveal well-formed, fully assembled 24-mer ferritin nanoparticles displaying uniformly distributed Env trimers.

**Figure S5.**
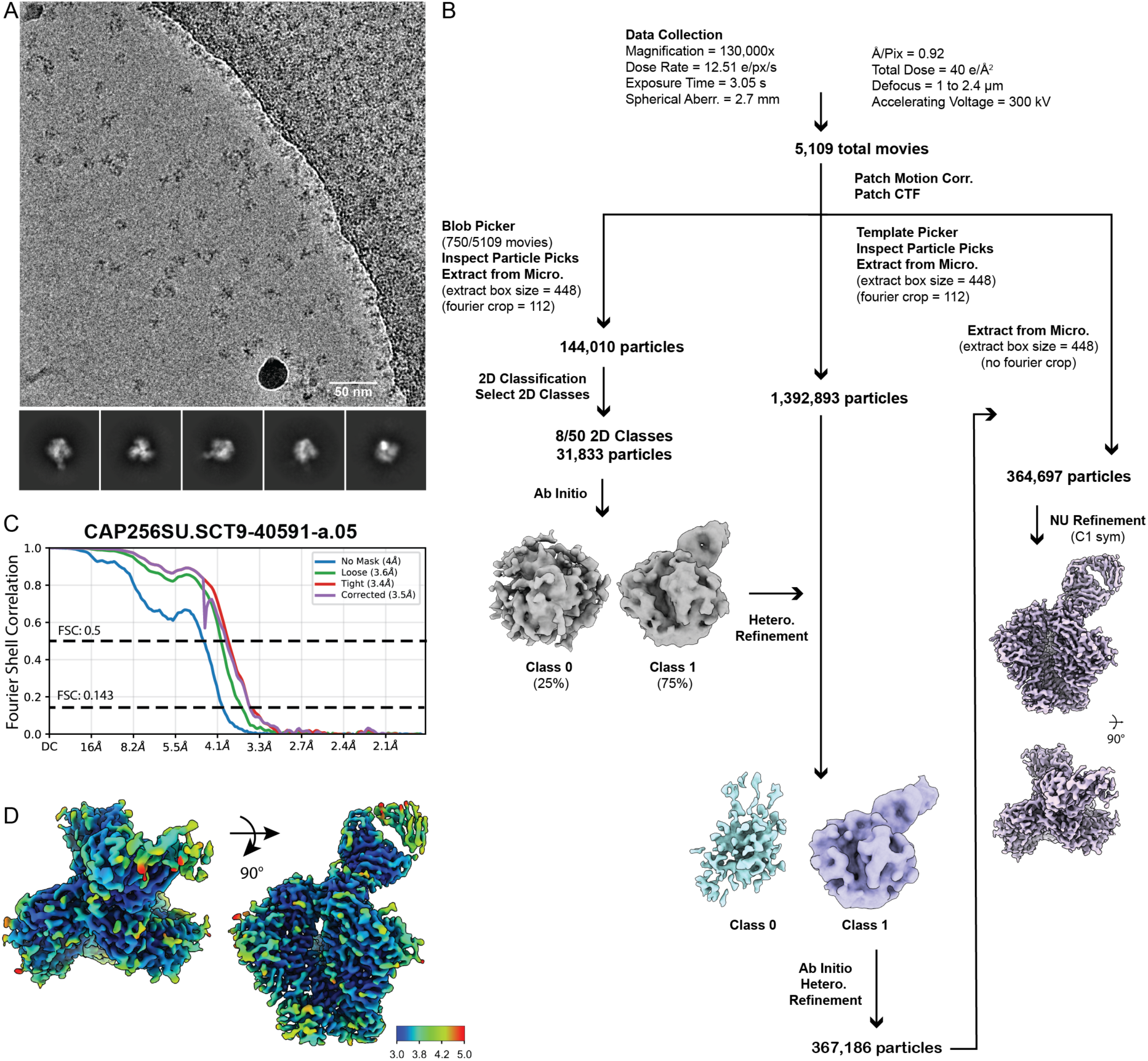
Cryo-EM data processing and validation for CAP256SU.SCT9 trimer in complex with 40591-a.05 Fab. **A.** Representative micrograph and representative 2D class averages for the CAP256SU.SCT9-40591-a.05 Fab complex. **B.** Workflow for single-particle cryo-EM data processing. Autopicking of a subset of motion-corrected and dose-weighted micrographs yielded 31,833 “good” particles that were used to generate an initial ab-initio model. 2D classes of these particles were used as templates to pick particles from the remaining movies in the dataset. Iterative rounds of ab initio and heterogeneous refinements on template-picked particles yielded 364,697 particles that correspond CAP256SU.SCT9 - 40519-a.05 Fab complex. These particles were refined to 3.5 Å resolution using C1 symmetry. **C, D.** Fourier shell correlation (FSC) plot (C) and local resolution estimation created in cryosSPARC **(D)** of the final reconstruction of the CAP256SU.SCT9 - 40591-a.05 Fab complex.

**Figure S6.**
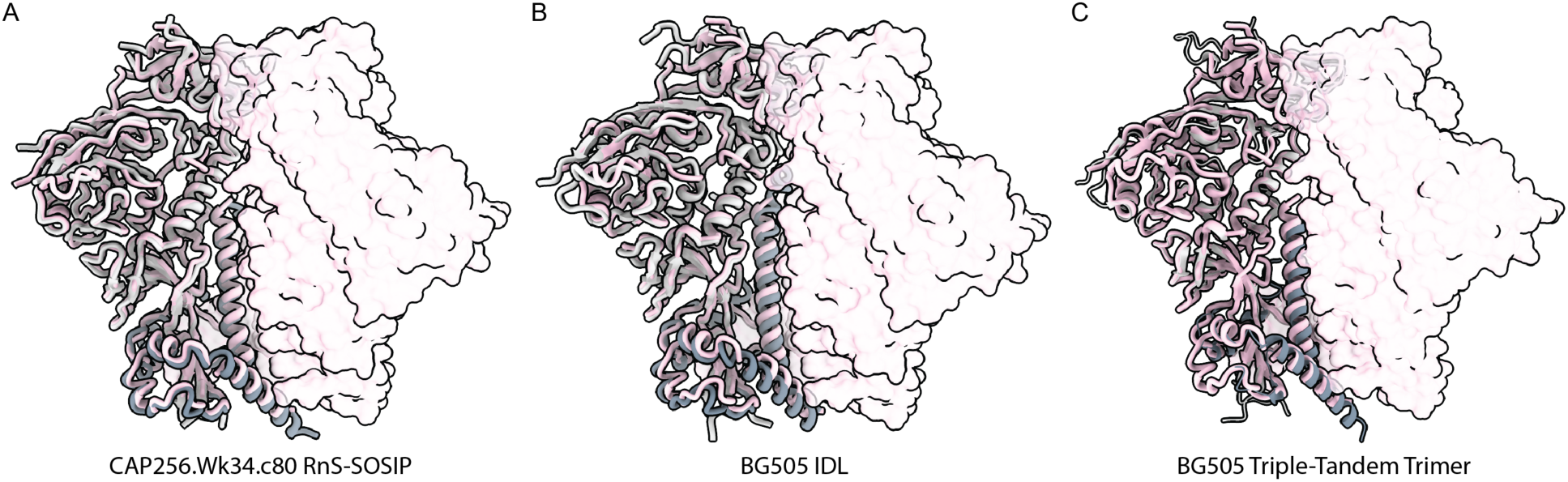
Comparisons of CAP256.SU-SCT9 global structure with alternate native-like Env stabilization strategies. Superposition of CAP256.SU-SCT9 trimer (shown in pink) with (**A**) CAP256.wk34.C80 RnS-SOSIP (shown in grey; PDB: 6VTT), (**B**) BG505 IDL (shown in grey; PDB 9BEW), and (**C**) BG505 triple-tandem trimer (shown in grey; PDB 6VTT). RMSD (0.88 Å, 0.974 Å, and 0.81 Å) calculations were determined from alignment of 387, 382, and 381 pruned Cα atoms pairs of gp140 subunit, respectively.

**Figure S7.**
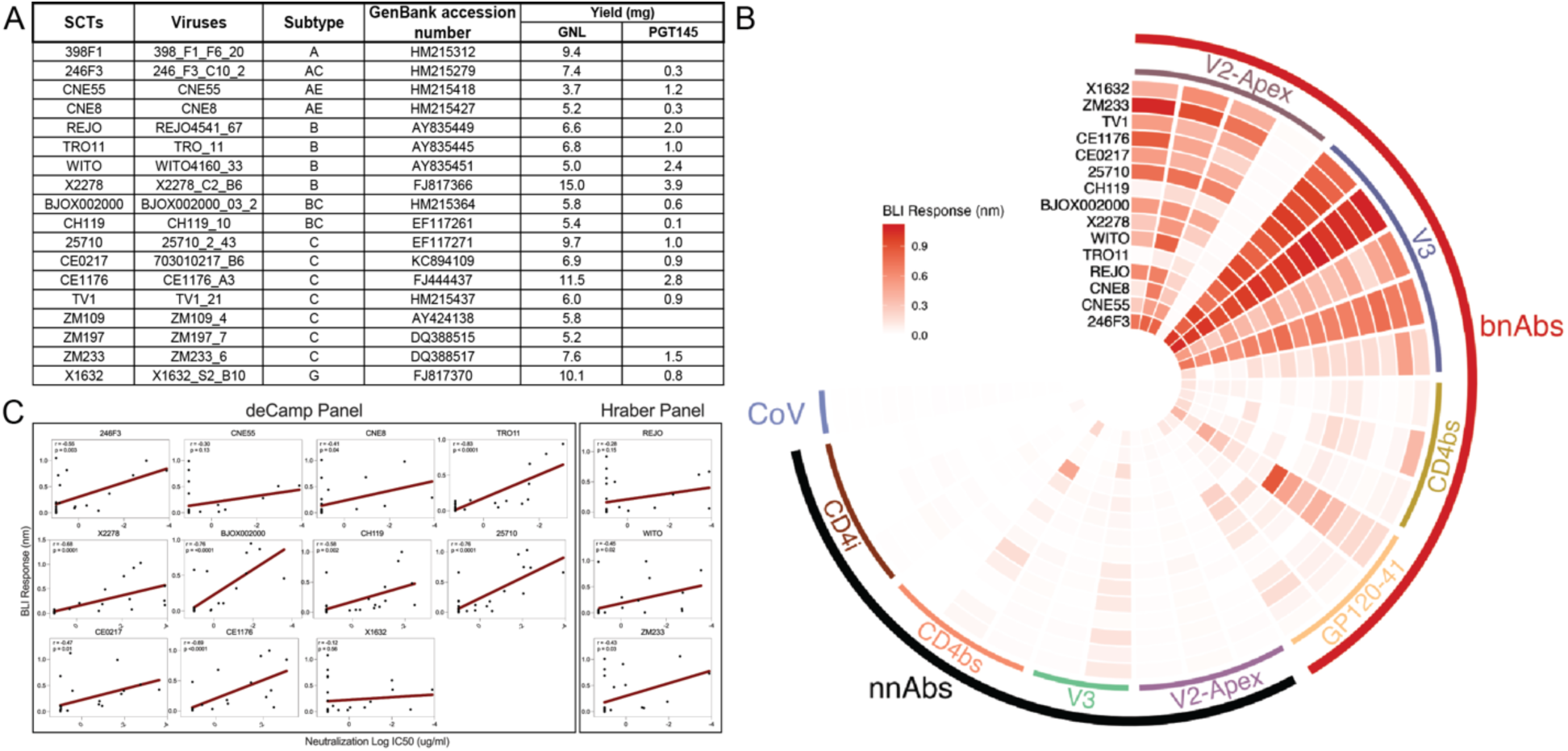
Antigenic characterization of PGT145 affinity-purified global HIV-1 panel of SCT-stabilized Env trimers. **A.** Table listing selected HIV-1 strains representing global genetic diversity. For each strain, the virus name, clade/subtype classification, and GenBank accession number are provided. Protein expression yields (mg/L) from transient 293F cell transfections are shown following purification by Galanthus nivalis lectin (GNL) and PGT145 antibody affinity chromatography. **B.** Heat map summarizing BioLayer Interferometry (BLI) maximum binding responses of PGT145-purified antibody affinity SCTs (100 nM) from the global panel to a select set of broadly bnAbs and non-neutralizing antibodies (nnAbs), each targeting specific epitopes on HIV-1 Env. Strong binding to bnAbs and minimal binding to nnAbs indicates that the SCTs maintain favorable antigenic profiles and present native-like conformational epitopes. **C.** Linear regression analysis (red line) showing a moderate negative correlation between BLI maximum binding responses and neutralization potency (log IC_50_ values).

**Figure S8.**
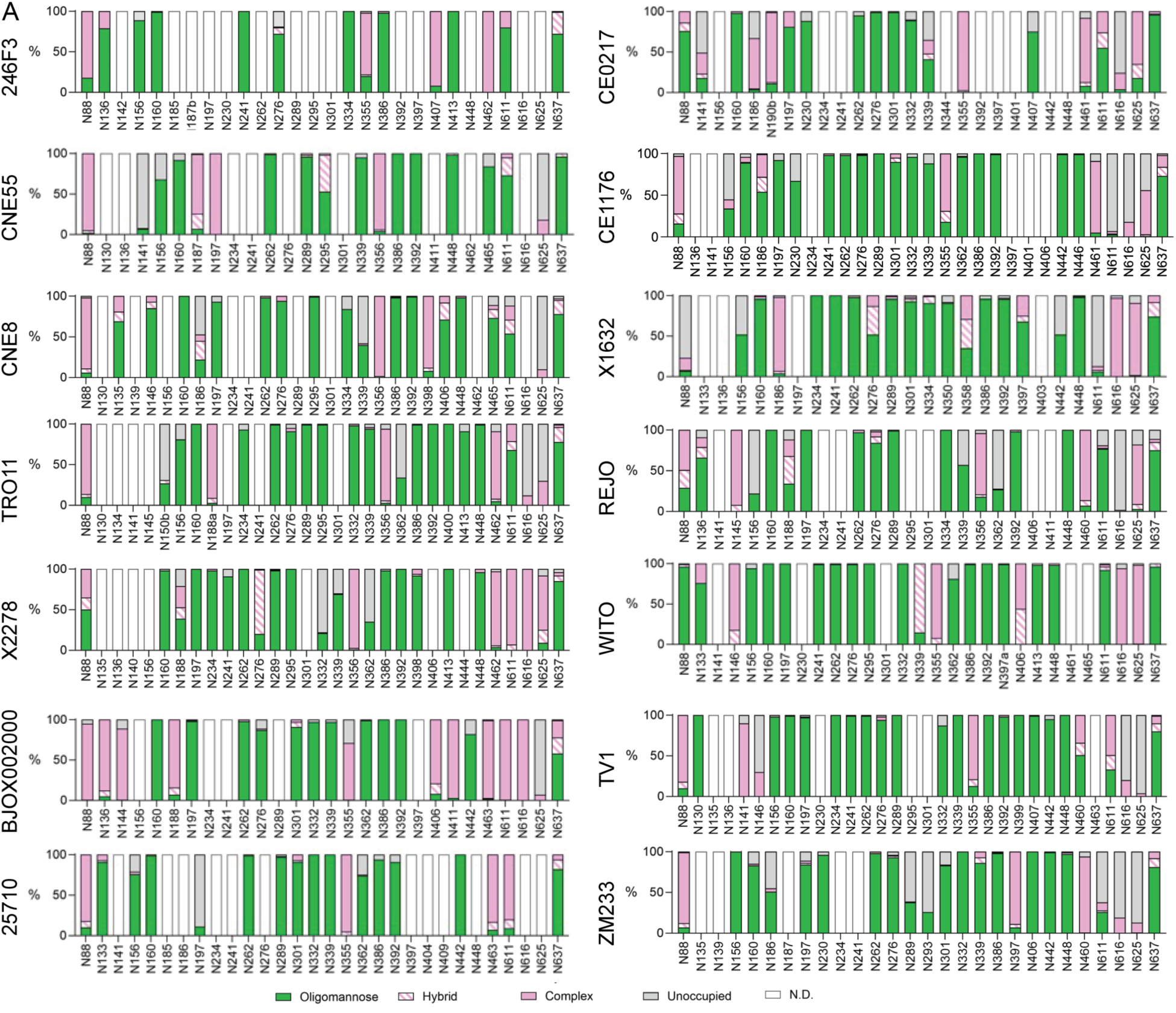
Site-specific glycan analysis of SCT-stabilized Env trimers from the global panel representing HIV-1 diversity. Bar plots summarize the glycan composition and site occupancy across the SCT-stabilized HIV-1 Env trimers representing diverse global subtypes. Each bar indicates the relative proportion of oligomannose-type, complex-type, hybrid-type, and unoccupied glycans at individual N-linked glycosylation (PNG) sites along the Env trimer. Site-specific glycan profiling was performed using mass spectrometry, enabling high-resolution mapping of glycosylation states across all analyzable sites. The data reveal that the SCTs exhibit a glycan landscape consistent with native-like HIV-1 Env, including high oligomannose occupancy at conserved sites commonly targeted by broadly neutralizing antibodies. These results support the structural integrity and immunological relevance of the SCT designs across globally diverse strains.

**Table S1.** Cryo-EM data collection and data processing statistics.

| CAP256SU.SCT9 - 40591-a.05 Fab |  |
| --- | --- |
| **Data Collection and Processing** |  |
| Microscope | Titan Krios |
| Camera | Falcon 4i |
| Magnification | 130,000x |
| Voltage (kV) | 30 |
| Recording Mode (?) | Counting |
| Dose Rate (e<sup>-</sup>/pixel/s) | 12.51 |
| Electron dose (e<sup>-</sup>/Å<sup>2</sup>) | 40 |
| Defocus Range (um) | -1 to -2.4 |
| Pixel size (Å) | 0.98 |
| Micrographs Collected | 11, 156 |
| Micrographs used | 5,109 |
| Total Extracted Particles | 1,392,893 |
| Refined Particles | 367,186 |
| **Reconstruction** |  |
| Final Particles | 364,697 |
| Symmetry Imposed | C1 |
| Nominal Resolution (Å) |  |
| FSC 0.143 (Unmasked/Masked) | 4/3.5 |
| Map Sharpening B-factor | 121 |
| **Refinement and Validation** |  |
| Number of Atoms | 16,823 |
| Protein | 2,039 |
| Ligand | 73 |
| MapCC | 0.82 |
| R.m.s. deviations |  |
| Bond lengths (Å) | 0.003 (0) |
| Bond angles (°) | 0.576 (0) |
| Clashscore (all atoms) | 8.45 |
| Poor rotamers (%) | 0 |
| Ramachandran plot |  |
| Favored (%) | 93.51 |
| Allowed (%) | 6.39 |
| Disallowed (%) | 0 |

**Table S2.**
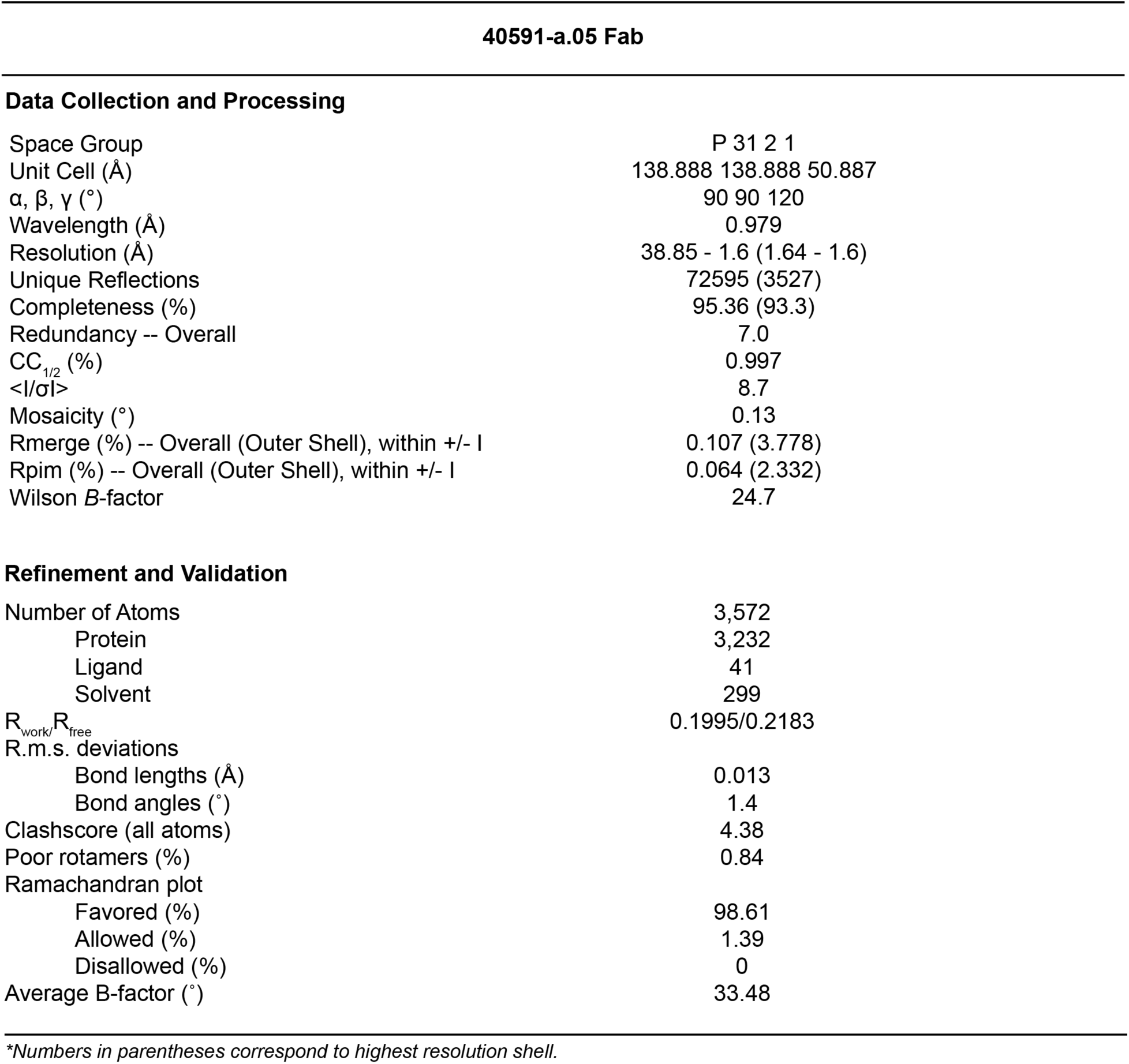
X-ray crystallography data collection and data processing statistics.

**Table S3.** Design and characterization of a large panel HIV-1 Envelope SCTs. Summary Table showing complete list of described SCTs, their associated GenBank accession number and the corresponding virus and accession numbers along with the subtypes. Protein yield/L 293F transfections through GNL affinity chromatography are shown.

| SCTs | Viruses | Subtype | GenBank accession number | Yield (mg) |
| --- | --- | --- | --- | --- |
| BG505 | BG505_W6M_C2 | A | DQ208458 | 6.0 |
| Q23 | Q23_17 | A | AF004885 | 3.9 |
| C1080 | C1080_C3 | AE | JN944660 | 7.0 |
| CM244 | CM244 | AE | AGL46598 | 15.0 |
| 40512 | 40512v26_env36 | AE | UMQ73315 | 7.1 |
| RV217 | RV217.40100 | AE | MN792078 | 3.8 |
| TH023 | TH023 | AE | AMB62180 | 5.2 |
| CRF250 | T250-4 | AG | MW507842 | 4.4 |
| 6563 | 6535.3 | B | AY835438 | 3.86 |
| B41 | B41 | B | EU576114 | 14.9 |
| 0646 | V703_0646_051_RE_con_s | C | UTT71827 | 4.93 |
| 2018 | H703_2018_240_RE_e6A1s | C | UTT72097 | 12.8 |
| 2117 | V703_2117_110_RE_sga2A10_s | C | UTT72103 | 24.5 |
| 2149 | H703_2149_060_RE_eB10s-VRC | C | UTT72109 | 10.2 |
| 2304 | V703_2304_150_RE_con_s | C | UTT72112 | 11.83 |
| 3728 | 3728_V2_C6 | C | HM215307 | 14.1 |
| 16055 | 16055_2_3 | C | EF117268 | 3.8 |
| 2759058 | 2759058_F10_B6 | C | HQ615959 | 7.2 |
| 2969249 | 2969249 | C | HQ595766 | 7.5 |
| 703010010 | 703010010_C4 | C | KC894105 | 6.4 |
| 704810053 | 704810053_2B7 | C | KC894119 | 6.4 |
| 0629 | H703_0629_150_RE_e5A2s | C | UTT71815 | 2.8 |
| B005582 | B005582_7_G7_8 | C | KF114882 | 5.6 |
| CAP256 | CAP256_SU | C | KF241776 | 5.6 |
| CAP382 | CAP382_2_00_D7_19 | C | KC154028 | 6.6 |
| CE2103 | CE2103_E8 | C | KC894086 | 6.3 |
| CH0694 | CH0694 | C | KJ700458 | 12.8 |
| CH505 | CH505_TF | C | KC247556 | 8.8 |
| CH848 | CH848_D1432_5_41 | C | KX217874 | 5.0 |
| Du156 | Du156.12 | C | DQ411852 | 6.8 |
| KO243 | KO243_H6_3 | C | KF114892 | 10.3 |
| S0431 | S0431_C1_1 | C | JN681246 | 5.3 |
| ZM215 | ZM215_8 | C | DQ422948 | 5.1 |
| C1086 | c1086 | C | FJ444395 | 7.4 |
| C1086_B2 | c1086 | C | FJ444395 | 6.6 |
| CH1012 | CH1012 | C | MG898887 | 5.3 |
| 191859 | 191859 | D | JX236672 | 9.1 |

